# Nasal microbionts differentially colonize and elicit cytokines in human nasal epithelial organoids

**DOI:** 10.1101/2024.09.25.614934

**Authors:** Andrea I. Boyd, Leah A. Kafer, Isabel F. Escapa, Amal Kambal, Hira Tariq, Susan G. Hilsenbeck, Hoa Nguyen-Phuc, Anubama Rajan, Joshua M. Lensmire, Kathryn A. Patras, Pedro A. Piedra, Sarah E. Blutt, Katherine P. Lemon

## Abstract

Nasal colonization by *Staphylococcus aureus* or *Streptococcus pneumoniae* is associated with an increased risk of infection by these pathobionts, whereas nasal colonization by *Dolosigranulum* species is associated with health. <u>H</u>uman <u>n</u>asal epithelial <u>o</u>rganoids (HNOs) physiologically recapitulate human nasal respiratory epithelium with a robust mucociliary blanket. We reproducibly monocolonized HNOs with these three bacteria for up to 48 hours with varying kinetics across species. HNOs tolerated bacterial monocolonization with localization of bacteria to the mucus layer and with minimal cytotoxicity compared to uncolonized HNOs. Human nasal epithelium exhibited both species-specific and general cytokine responses, without induction of type I interferons, consistent with colonization rather than infection. Only live *S. aureus* colonization robustly induced IL-1 family cytokines, suggestive of inflammasome signaling. *D. pigrum* and live *S. aureus* decreased CXCL10, whereas *S. pneumoniae* increased CXCL11, chemokines involved in antimicrobial responses to both viruses and bacteria. Overall, HNOs are a compelling model system to reveal host-microbe dynamics at the human nasal mucosa.

**IMPORTANCE:** Human nasal microbiota often includes highly pathogenic members, many of which are antimicrobial resistance threats, e.g., methicillin-resistant *Staphylococcus aureus* and antibiotic-resistant *Streptococcus pneumoniae*. Preventing colonization by nasal pathobionts decreases infections and transmission. In contrast, nasal microbiome studies identify candidate beneficial bacteria that might resist pathobiont colonization, e.g., *Dolosigranulum pigrum*. Discovering how these microbionts colonize the human nasal passages and means to reduce pathobiont colonization is limited by previous models. This creates an urgent need for human-based models that exemplify bacterial nasal colonization. We addressed this need by developing human nasal epithelial organoids (HNOs) as a new model system of bacterial nasal colonization. HNOs accurately represent the mucosal surface of the human nasal passages enabling exploration of bacterial-epithelial interactions, which is crucial since the epithelium instigates the initial innate immune response to bacteria. Here, we identified differential epithelial cytokine responses to these three bacteria setting the stage for future research.

## INTRODUCTION

Methicillin-resistant *Staphylococcus aureus* (MRSA) is a leading cause of death due to antimicrobial resistant bacteria globally (1). The human nasal passages are a primary habitat for *S. aureus* with a third of humans nasally colonized (2, 3). Moreover, *S. aureus* nasal colonization is a risk factor for *S. aureus* infection with ∼80% of infection isolates matching the person’s nasal isolate (4–6). In the absence of an effective vaccine (7, 8), antibiotic-based *S. aureus* nasal decolonization is the only current strategy to reduce infections (9–11). Further, evolutionary analysis shows humans are the major hub for *S. aureus* host switching among mammalian species (12). Thus, development of human-based models to investigate *S. aureus* nasal colonization is vitally important for the World Health Organizations’ One Health, which is “an integrated, unifying approach that aims to sustainably balance and optimize the health of people, animals, and ecosystems” (https://www.who.int/health-topics/one-health#tab=tab_1). Similarly, *S. pneumoniae* human nasal colonization is the primary reservoir for both invasive pneumococcal disease (IPD) and for pneumococcal transmission (13). Vaccination has reduced IPD; however, in the absence of a universal vaccine for *S. pneumoniae*, IPD remains a global threat to human health (1, 14). To date, experimental human pneumococcal colonization studies are limited to only a few serotypes (15), and it is unknown whether findings are applicable to a broader range of serotypes, including highly virulent serotypes. In contrast to the threat posed by these two nasal pathobionts, the genus *Dolosigranulum* is frequently associated with healthy cohorts in studies of human nasal microbiota (16) and is inversely associated with *S. aureus* nasal colonization in adults (17–19) and with *S. pneumoniae* nasal colonization in children (20, 21). Furthermore*, D. pigrum* inhibits *S. aureus in vitro* (18) and protects *Galleria mellonella* from *S. aureus* infection *in vivo* (22), supporting its likely role as a nasal mutualist. Using these three species, we demonstrate here that human nasal epithelial organoids (HNOs; aka human nose organoids) are a physiologically relevant model system for elucidating conserved and species-specific human microbiont-epithelial interactions.

HNOs offer more physiological accuracy, genetic diversity, and long-term experimental use than other colonization models. Based on the Sachs *et al*. method for generating Human Airway (bronchial) Organoids (23), HNOs are derived from tissue-resident stem cells collected using nasal mucosal swabs/washings and propagated *ex vivo* as three-dimensional (3D) organoids that can be frozen and passaged for long-term experimental use. Dispersed 3D organoids plated in monolayers on transwells and differentiated at an air-liquid interface (ALI) accurately recapitulate human nasal respiratory epithelium with ciliated cells, goblet cells, basal cells, club cells, and a thick mucus layer that is circulated by functional cilia and in contact with air (24, 25). HNOs differentiated at ALI have been used as a model system for respiratory viral infections (24–27); however, this model has not been previously applied to study host-bacterial interactions in the human nasal passages.

HNOs differentiated at ALI fill crucial gaps between existing nasal colonization models and overcome many limitations of other model systems for investigating epithelial-microbiont interactions. Primary human nasal epithelial cell explants (pHNECs), although physiologically equivalent, are restricted to short-term studies due to limited passaging (28). Immortalized respiratory cell lines (e.g., Calu-3, RPMI 2650, A549 cells) lack a physiological apical mucus layer (although Calu-3 cells do produce mucus), lack the multiple cell types present in human nasal respiratory epithelium, and fail to represent human genetic diversity (29, 30). ALI-polarization of these cell lines (31–33) or of immortalized human nasal epithelial cells (34) generates models that can tolerate live bacterial colonization for 4 to 72 hours, depending on the species (31–34); however, these still fall short of recapitulating the human nasal mucociliary blanket. Animal models, e.g., the cotton rat for *S. aureus* (35) and infant mice for *S. pneumoniae* (36), have the advantage of a systemic immune response; however, many human nasal microbionts poorly colonize nonhuman models and animal models of human microbiota may nominally reflect human-microbe interactions (37). The data presented here demonstrate that HNOs overcome these limitations providing a new model system for studying bacterial-epithelial interactions for common members of the human nasal microbiota.

## RESULTS

### Human nasal epithelial organoids differentiated at air-liquid interface produce a robust apical mucus layer

Fixing HNOs differentiated at ALI in Clark’s solution (38, 39) and staining with Periodic acid-Schiff (PAS) and hematoxylin revealed a substantial apical mucus layer atop the epithelium with visible apical ciliated cells, goblet cells, and numerous basal cells (**Fig. 1A**). This robust mucus production by HNOs differentiated at ALI is a vital element since mucus affects bacterial behavior by reducing adherence and biofilm formation (40, 41). Henceforth, we refer to HNOs differentiated at ALI simply as HNOs. To replicate *in vivo* environmental conditions, we performed experiments with HNOs at the human nasal passage temperature of 34 °C (42). Accounting for human nasal temperature is particularly important since *S. aureus* (33, 43–45) and *S. pneumoniae* (46, 47) behave differently at 34 °C, or lower, compared to their behavior at the human internal body temperature of 37 °C. Based on bulk RNA sequencing, HNOs differentiated at 37 °C and shifted to 34 °C for experimentation were transcriptionally similar to those maintained at 37 °C throughout (**Fig. S1A**), indicating the shift to 34 °C had minimal effects on epithelial transcription. Hypothesizing that we could use these HNOs to investigate bacterial nasal colonization, we developed protocols for live bacterial monocolonization by *S. aureus*, *S. pneumoniae,* and *D. pigrum*. Of note, as shown in Figure 1, we define colonization as the bacteria residing in the mucus layer without microscopic evidence of epithelial damage and with a low-level release of cytoplasmic lactate dehydrogenase (LDH, a damage indicator) similar to what occurs in uncolonized HNOs. This contrasts with infection, which is defined as the bacteria causing substantial damage to the HNO within 24 hours.

**Figure 1.**
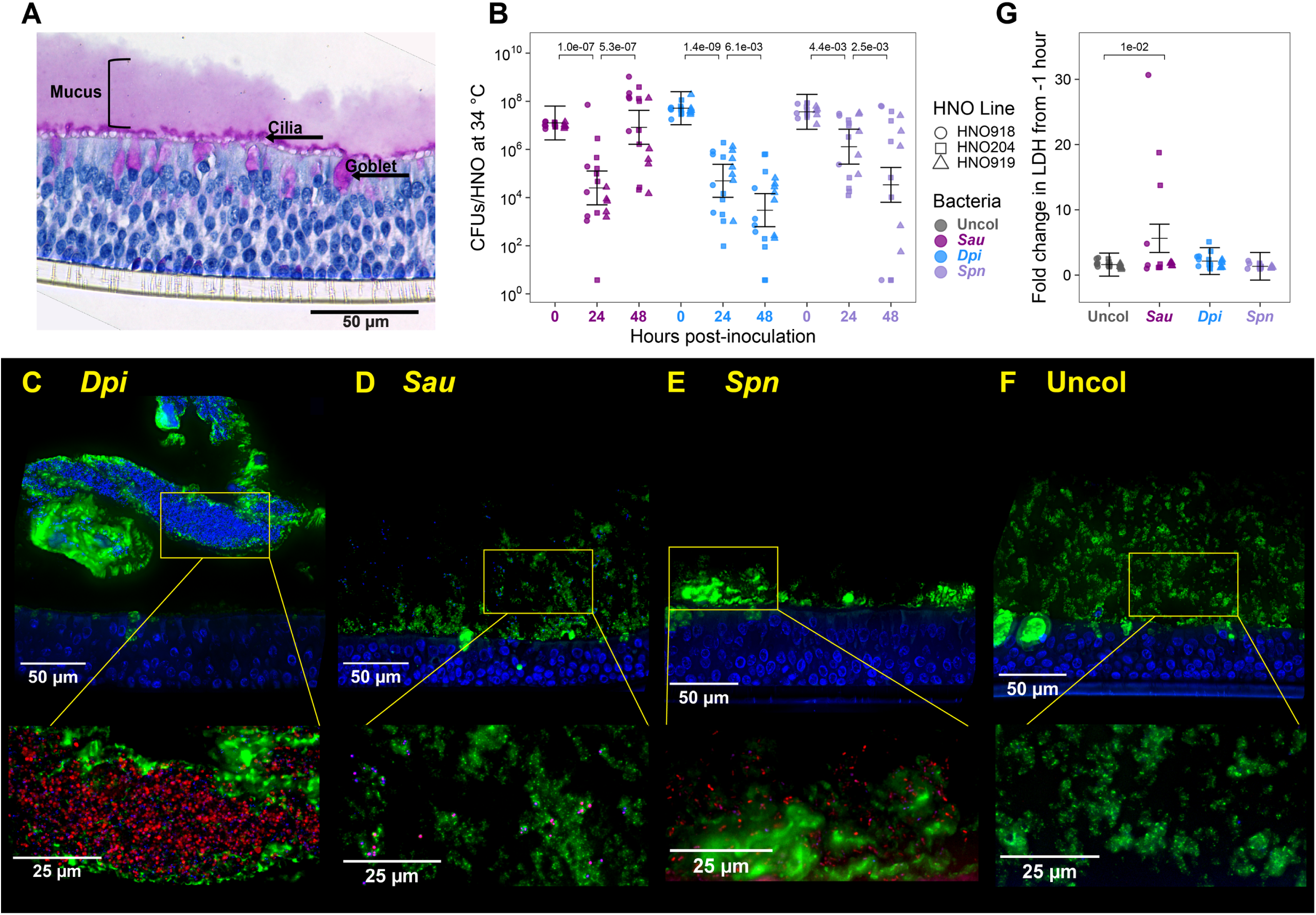
Human nasal epithelial organoids tolerate nasal microbiont colonization and restrict bacteria to the mucus layer. (**A**) HNOs differentiated at an ALI produced a robust mucus layer (pink). Uncolonized HNO919 was fixed in Clarke’s solution, cross-sectioned, and stained with PAS (magenta) to highlight goblet cells, secreted mucus, and cilia, and with hematoxylin (navy) to highlight epithelial cells then imaged at 40x magnification. (**B**) Three HNO lines (HNO918 circles, HNO204 squares, HNO919 triangles) were each monocolonized with *S. aureus* (purple), *D. pigrum* (blue), and *S. pneumoniae* (lavender) at 34 °C for up to 48 h. At time 0, 10^7^ CFUs of a bacterium in 15 µL of EBSS were inoculated apically. Recovered CFUs/HNO at 24 and 48 h are shown. Independent experiments for B and G: HNO918 ≥ 3, HNO204 ≥ 7, and HNO919 ≥ 5. Data (B and G) were analyzed using a linear mixed-effects model (LMM) to determine statistical significance with *p*-values adjusted for multiple comparisons, shown above the horizontal bars (0 to 24 h and 24 to 48 h in B and Uncolonized to each bacterial species in G). Vertical brackets represent the model-predicted mean values and confidence intervals (+/- twice the standard error of the mean). Including HNO line in the LMM as a random effect showed that HNO line accounted for ≤ 2.4% of the variance (**Table S1A-B**). (**C**) *D. pigrum,* (**D**) *S. aureus*, and (**E**) *S. pneumoniae* localized in the mucus layer at 6 h of HNO monocolonization. (**F**) Uncolonized HNO control. Shown are representative fluorescent images of cross-sections of colonized or uncolonized HNOs fixed and stained with anti-MUC5AC antibody highlighting mucus and goblet cells (green), and Hoechst highlighting host nuclei and bacterial cells (blue) at 60x magnification. Yellow boxes indicate the area that is shown below at a higher magnification (100x) to highlight bacteria (prestained with MitoTracker Red CMXRos prior to colonization) in the mucus layer. (**C**-**F**) Representative images are from experiments done with lines HNO204 and HNO918, each assayed on a different day: (**C**) HNO918, (**D**) HNO204, (**E**) HNO918, (**F**) HNO204. The other images are at https://github.com/KLemonLab/HNOBac_Manuscript/tree/main/data/microscopy. (**G**) Fold change in lactate dehydrogenase (LDH) release of uncolonized (gray) HNOs and of HNOs colonized with *S. aureus* (purple), *D. pigrum* (blue), or *S. pneumoniae* (lavender) in HNO basal medium at 48 h compared to −1 h samples from the same well.

### Human nasal pathobionts and a mutualist monocolonize HNOs

All three nasal microbionts monocolonized HNOs at 34 °C, as measured by colony forming units (CFUs) recovered per HNO at 24 and 48 hours (**Fig. 1B**). To preserve the mucus layer and maintain the air interface, we apically inoculated HNOs with 10^7^ - 10^8^ CFUs of bacteria in only 15 µL of buffer, with control HNOs receiving buffer alone. To account for host genetic variability, we used three different donor-derived HNO lines, which are indicated by different shapes in graphs. We used a linear mixed-effects model (LMM) to account for the hierarchical nature of our data, with n ≥ 3 independent experiments within each HNO line. The LMM allowed us to assess the variance due to both HNO line and independent experiments as random effects. The variance due to HNO line was minimal when modeling the CFU and LDH data (**Table S1A-B**), accounting for 0-to- 2.8% of the variance in Figures 1B, 1G, S1C, and S1D. This allowed us to appropriately group the different HNO lines together in each of these graphs and to focus the model on the different bacterial treatments as a fixed effect. (For details on statistical analysis see reproducible code available at (https://klemonlab.github.io/HNOBac_Manuscript).

The 3 HNO donor lines had a median of 6.68 x 10^5^ human cells per transwell one day prior to bacterial inoculation, based on counts from a subset of the experiments in this study (**Fig. S1B**). Using the median number of cells per transwell for each HNO line, the estimated range of the multiplicity of inoculation for all experiments was 14 – 65. At 24 hours, *S. aureus* CFUs/HNO were reduced 500-fold from the inoculum, with an LMM-predicted mean of ∼2.5 x 10^4^ (**Table S1A**), but then appeared to rebound, increasing 330-fold between 24 and 48 hours (purple, **Fig. 1B**), indicating replication on the mucosal surface. We modeled *S. aureus* nasal colonization using an epidemic USA300 MRSA strain because of its clinical importance in causing invasive infection (48). At 24 hours, *D. pigrum* CFUs/HNO were reduced 1043-fold from the inoculum, with a model-predicted mean of ∼5.0 x 10^4^ CFUs/HNO, which further decreased 17-fold by 48 hours (blue, **Fig. 1B**). At 24 hours of colonization, *S. pneumoniae* CFUs/HNO were reduced 28-fold from the inoculum, with a model-predicted mean of ∼1.3 x 10^6^ CFUs/HNO, which further decreased 39-fold by 48 hours (lavender, **Fig. 1B**). Experiments at human internal body temperature of 37 °C gave similar CFU results to those at 34 °C for each bacterium (**Fig. S1C**), based on statistical analysis comparing each bacterium at a given time point between both temperatures (**Table S1A**). Together, these assays demonstrate persistence of close to 10^5^ CFUs/HNO at 24 or 48 hours, providing theoretically sufficient bacterial biomass for downstream assays in future experimentation. Moreover, these data point to the ability of the nasal epithelium to control the bacterial biomass during colonization.

### Bacteria reside in the mucus layer during colonization

To assess bacterial localization, we colonized with live bacteria that were prelabeled with MitoTracker Red CMXRos (49), which has fluorescence that survives fixation. Then, after HNO fixation and sectioning, we visualized the mucus layer and goblet cells using an anti-MUC5AC antibody and the epithelial cell nuclei and bacterial DNA using Hoechst 33342 dye (**Fig. 1C-E**). Both colonized HNOs and uncolonized controls (**Fig. 1F**) showed apical secretion of MUC5AC-positive mucus. Moreover, bacteria were visible in the apical mucus layer of HNOs after 6 hours of colonization (**Fig. 1C-E**). This supports prior reports of *S. aureus* localization in mucus during nasal colonization of ferrets (50) and of *S. pneumoniae* localization in mucus layer during colonization of nasal epithelial tissue explants (51). We observed variability in how well the MitoTracker Red CMXRos appeared to label each species (best for *D. pigrum*), which is a limitation of this approach.

### HNOs tolerate bacterial colonization

To assess HNOs for cellular damage following bacterial colonization, we measured lactate dehydrogenase (LDH), an enzyme that is released from epithelial cells when plasma membrane integrity is compromised, in the basal medium (52). For uncolonized HNOs, LDH levels in the basal medium increased 1.6-fold between 1 hour before colonization to 48 hours (**Fig. 1G**). HNOs colonized with *D. pigrum* or *S. pneumoniae* released similar amounts of LDH as uncolonized controls (**Fig. 1G, Table S1B**). HNOs colonized with *S. aureus* released slightly (3.5-fold) more LDH by 48 hours than did uncolonized controls (**Fig. 1G, Table S1B**). Similar results were obtained at 37 °C for each bacterium (**Fig. S1D**), based on statistical analysis comparing LDH results for each bacterium between both temperatures (**Table S1B**). These data indicate a similarly low level of cellular damage in uncolonized and bacterially colonized HNOs.

### HNOs display characteristics of human nasal respiratory epithelium that are absent in Calu-3 and RPMI 2650 cells

HNOs produce the cell types expected of human nasal respiratory epithelium (24, 25). Based on this, we hypothesized that they would display characteristics of human nasal respiratory epithelium that are lacking in cancer-derived cell lines commonly used to model bacterial nasal colonization, such as Calu-3 and RPMI 2650. Calu-3 cells were isolated in 1975 from a metastatic pleural effusion in a 25-year-old man with adenocarcinoma of the lung (53). RPMI 2650 cells were isolated in 1962 from a metastatic pleural effusion in a 52-year-old-man with an anaplastic squamous cell carcinoma of the nasal septum (54). To test our hypothesis, we compared uncolonized HNOs differentiated at ALI to Calu-3 and RPMI 2650 cells polarized at ALI using brightfield and immunofluorescence microscopy (**Fig. 2**). Each epithelial model (in transwells) was fixed in Clark’s solution (38, 39) prior to cross-sectioning and staining. PAS and hematoxylin staining again showed that HNOs have a thick apical mucus layer, along with ciliated apical cells and goblet cells (magenta in **Fig. 2A**), as was shown in Figure 1A. In contrast, Calu-3 cells had scant apical mucus with most of the mucus visible as pockets within the cell layer (**Fig. 2B**) and RPMI 2650 cells lacked any visible mucus (**Fig. 2C**). The mucus findings were the same using IFM with an anti-MUC5AC antibody (**Fig. 2D-F**), which better highlights the pockets of mucus intercalated between the Calu-3 cells (**Fig. 2E**). These findings are consistent with prior reports for Calu-3 (55, 56) and RPMI 2650 cells (57, 58). Of the three models, only HNOs displayed alpha-tubulin-positive apical ciliated cells and KRT5-positive basal cells (Fig. 2G). Based on these findings, HNOs fill a crucial gap as a model system because they have characteristics of human nasal respiratory epithelium that are absent in cancer-derived cell lines.

**Figure 2.**
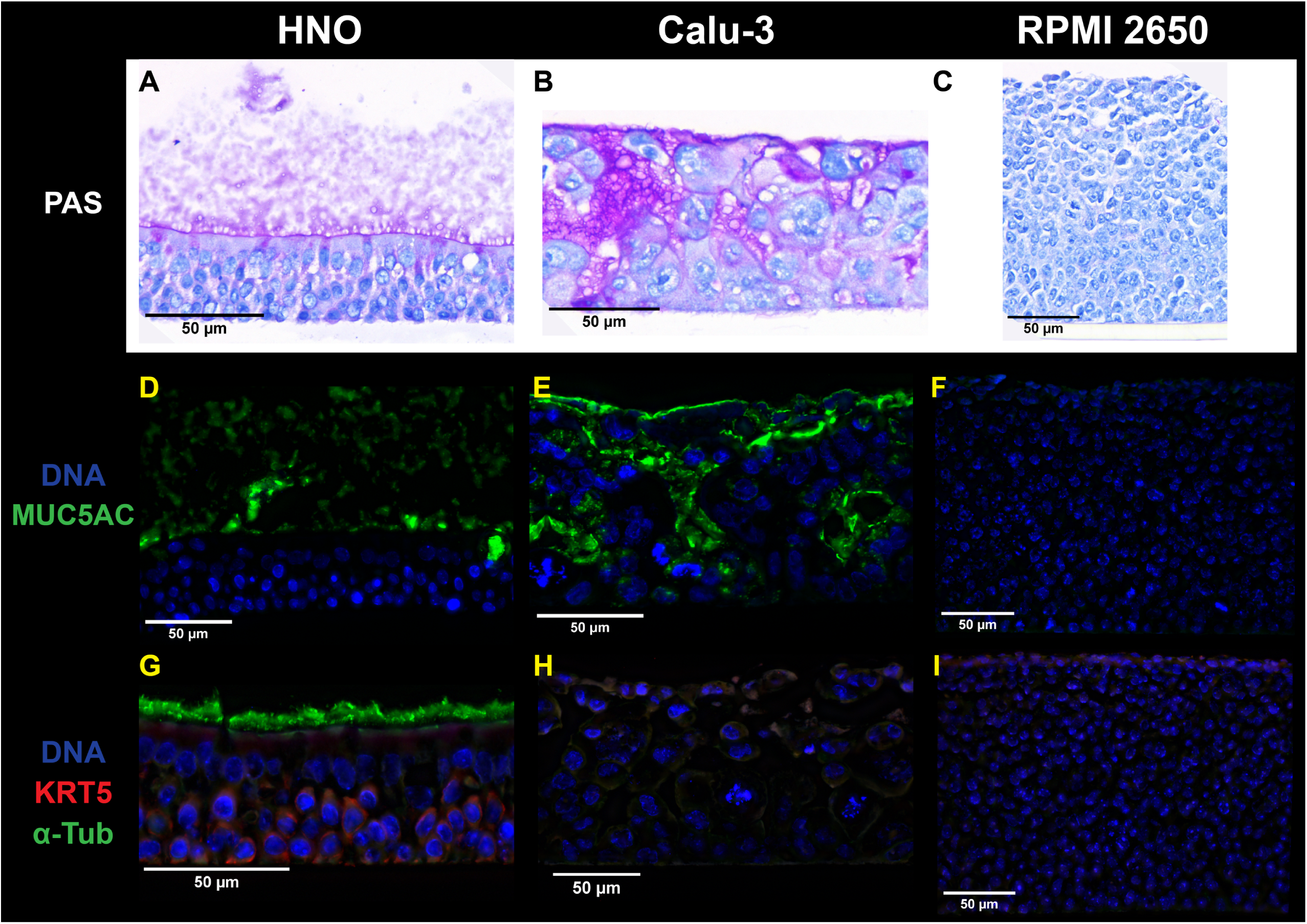
HNOs exhibit many characteristics of human nasal respiratory epithelium that are lacking in Calu-3 and RPMI 2650 cells. Representative brightfield microscopy images of uncolonized (**A**) HNO204, (**B**) Calu-3, and (**C**) RPMI 2650 cells at ALI stained with PAS (magenta) to highlight goblet cells, secreted mucus, and cilia and hematoxylin (navy) to highlight nuclei. Representative immunofluorescence microscopy images from uncolonized (**D**) HNO204, (**E**) Calu-3, and (**F**) RPMI 2650 cells stained with anti-MUC5AC antibody (green) to highlight mucus and goblet cells and Hoechst (blue) to highlight nuclei. Adjacent sections from the same samples were stained with (**G-I**) anti-KRT5 antibody (red) to highlight basal cells, anti-acetylated alpha-tubulin antibody (green) to highlight cilia, and Hoechst (blue) to highlight nuclei (**D-I**). Images are cropped to display the full thickness of the epithelium plus the mucus layer (for HNOs) such that the final images are not sized proportionally across the three models (as reflected by the scale bars). All cells at ALI in transwells were fixed in Clarke’s solution prior to cross sectioning. Images were acquired at a magnification of 40x (**A-C**) or 60x (**D-I**). Images are representative of multiple independent experiments: *n* = 3 in HNO204, *n* = 3 in Calu-3 cells, and *n* = 2 in RPMI 2650. Additional images are at https://github.com/KLemonLab/HNOBac_Manuscript/tree/main/data/microscopy.

In contrast to the stark differences in mucus production and the cell types present (**Fig. 2**), all three epithelial models (HNOs, Calu-3, and RPMI 2650) supported comparable levels of bacterial monocolonization at 6 hours, as reflected by the log2-fold change in CFUs compared to the inoculum for *S. aureus*, *D. pigrum*, and *S. pneumoniae* (**Fig. 3A**). Similarly, 6 hours of bacterial monocolonization resulted in comparably low levels of basal LDH release (indicative of minimal cell damage) across all three models for each bacterium (**Fig. 3B**). We compared the three models at 6 hours based on reports that polarized Calu-3 cells would tolerate USA300 *S. aureus* monocolonization in this time frame (31).

**Figure 3.**
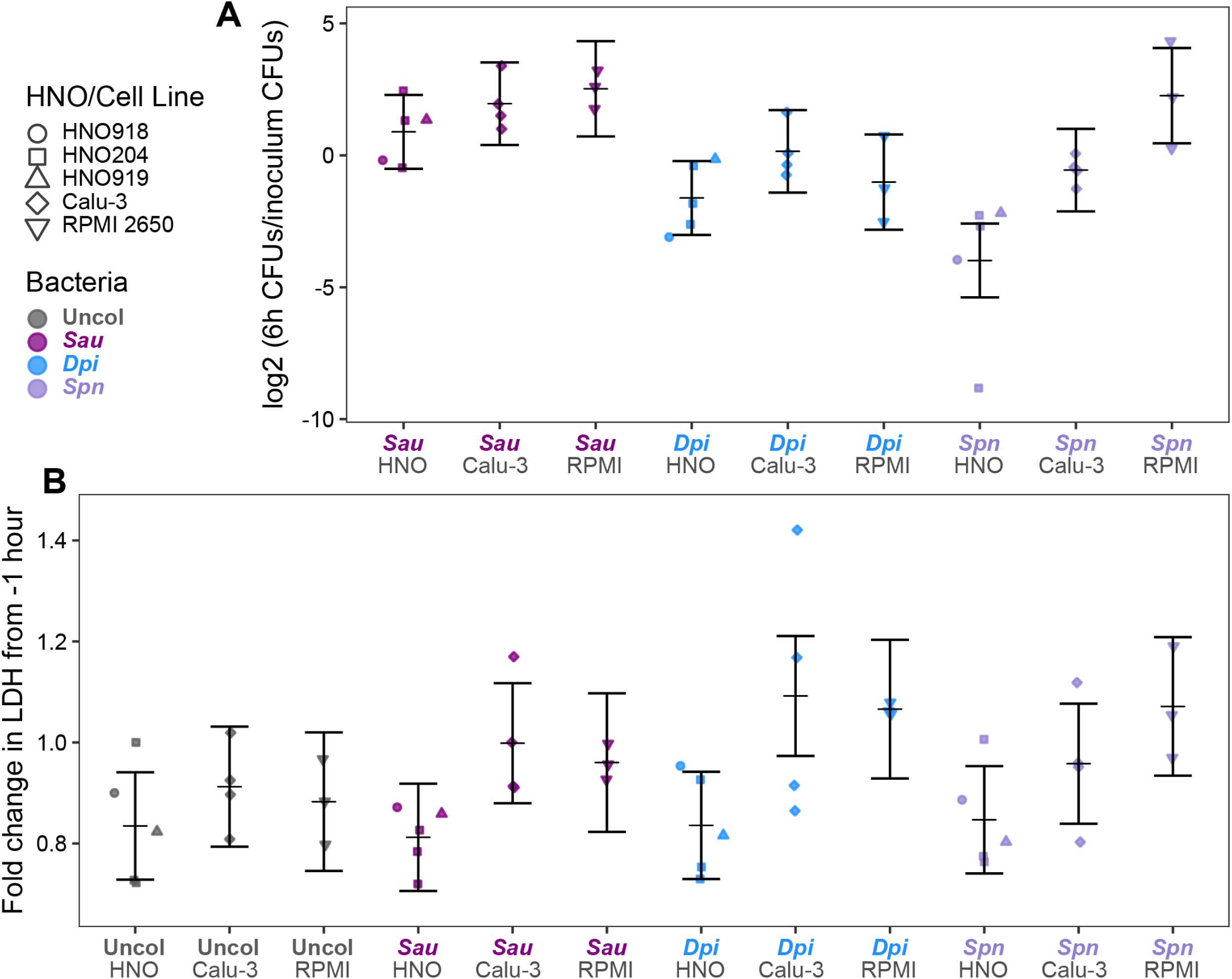
HNOs, Calu-3 cells, and RPMI 2650 cells at ALI exhibit similar levels of bacterial colonization and epithelial cell damage at 6 hours. Differentiated HNOs (circle, square, triangle), polarized Calu-3 cells (diamond), and polarized RPMI 2650 cells (upside-down triangle) at ALI all (**A**) supported comparable levels of CFUs (based on the log2-fold change of CFUs at 6 h compared to the inoculum) and (**B**) produced comparable levels of basal LDH (based on fold change compared to 1 h before inoculation) when monocolonized with each bacterial species for 6 hours. *S. aureus* (purple), *D. pigrum* (blue), *S. pneumoniae* (lavender), or uncolonized (gray). No statistically significant differences were detected using a LMM with comparisons between all pairs of epithelial models within each tested bacterial species (**Table S1A-B**). The vertical brackets represent the model-predicted mean values and confidence intervals (+/- twice the standard error of the mean). Independent experiments: *n* = 5 for HNOs (HNO204 = 3, HNO918 = 1, HNO919 = 1); *n* = 4 for Calu-3; *n* = 3 for RPMI 2650.

### HNOs display both species-specific and general cytokine production in response to monocolonization with *S. aureus*, *D. pigrum*, or *S. pneumoniae*

Based on their pathogenic potential, we hypothesized the pathobionts *S. aureus* and *S. pneumoniae* would initiate a more prominent nasal epithelial innate immune response compared to the candidate mutualist *D. pigrum*. To test this, we measured HNO apical and basal cytokine production in response to monocolonization with each species, with apical production modeling release into human nasal mucus/lumen and basal production modeling release into tissue and circulation. We assayed a wide range of epithelial-produced cytokines (including those involved in allergic response, inflammasome signaling, matrix metalloproteinases, acute inflammation, chemotaxis, growth factors, response to viral/bacterial infection, and anti-inflammation), totaling 39 cytokines, 30 of which met the threshold for detection in at least one compartment (apical and/or basal). To assess whether live and dead bacteria elicited similar effects, we also inoculated HNOs with a comparable number of dead bacteria killed by gamma irradiation, which should preserve the structure of surface-exposed antigens (59). We visualized the log_2_-fold-change in each cytokine detected at 48 hours in each colonized condition vs. the uncolonized control HNO, which revealed both species-specific and general responses to bacterial colonization (**Fig. 4; Table S1C**).

**Figure 4.**
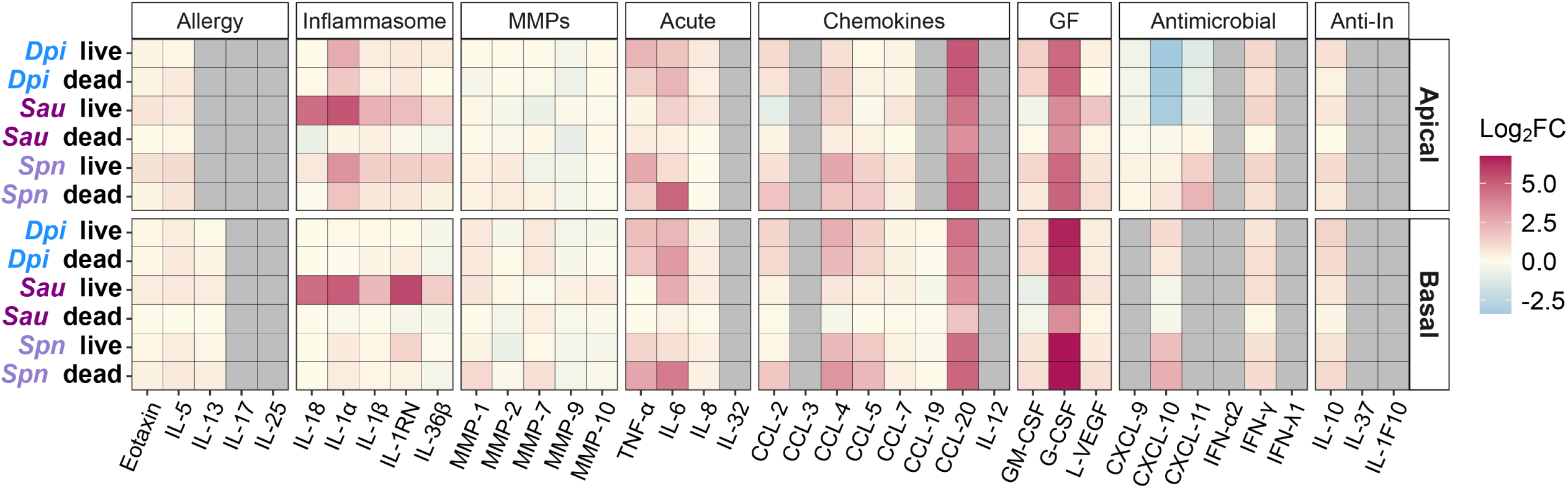
The human nasal respiratory epithelium of HNOs produce general and species-specific cytokine responses to bacterial colonization. A heatmap shows the log2-fold change of the ratio between the average amount of each of 39 cytokines assayed for in an apical wash or in the basal medium of HNOs monocolonized with live or dead (gamma-irradiated) bacteria compared to the uncolonized control at 48 h of colonization at 34 °C (**Table S1C**). Cytokines are grouped by primary epithelial function. Gray tiles indicate levels below our threshold for limit of detection (see Methods). IL-12 is IL-12p70. MMPs are matrix metalloproteinases. GF is growth factors. Anti-In is anti-inflammatory.

In terms of a species-specific effect, *S. aureus* colonization induced interleukin-1 (IL-1) family cytokines consistent with an inflammasome response (**Fig. 5A**). Inflammasomes are cytosolic multiprotein complexes that are assembled in response to pathogen-associated molecular patterns (PAMPs) and/or damage-associated molecular patterns (DAMPs). Inflammasomes serve as both sensors and effectors of downstream inflammatory responses usually leading to pyroptotic cell death, which releases intracellular defenses (60) and DAMPs that enhance recruitment of immune cells to the site of infection (61). Monocolonization with live *S. aureus* increased release of interleukin-1 alpha (IL-1α) by 39-fold apically and 33-fold basally compared to uncolonized HNOs (**Fig. 5A**). Again, we used an LMM for analysis, which showed that HNO line accounted for little of the variance (**Table S1D**). The fold changes mentioned here are based on the LMM predicted means (**Table S1E**). IL-1α is a key alarmin initiating inflammasome signaling (62). Both the pro- and mature forms of IL-1α bind to and actively signal through the IL-1 receptor (IL-1R) (63), and the Luminex assay used here detects both forms. In opposition, IL-1 receptor antagonist (IL-1RN, previously known as IL-1RA) competitively binds to the IL-1R inhibiting both the pro- and mature forms of IL-1α and dampening inflammasome activation. Live *S. aureus* monocolonization also increased HNO production of IL-1RN by 4-fold apically and 50-fold basally. This raised the question of whether IL-1α or IL-1RN is dominant in the HNO response to *S. aureus* colonization. Using the same samples, we detected net IL-1R activation in the HEK-Blue IL-1R reporter line indicating that IL-1α activity dominated apically (**Fig. S2A**) and basally (**Fig. S2B**) during live *S. aureus* colonization. Cells undergoing stress or damage release IL-1α, leading to inflammasome activation and subsequent release of mature IL-18 and/or IL-1β (62). In response to live *S. aureus* colonization, HNOs also released higher levels of IL-18 (but not IL-1β) with a 23-fold apical and basal increase, suggesting inflammasome activation in a subset of epithelial cells in line with a very modest increase in LDH release under this condition (**Fig. 1G**). In contrast, gamma-irradiated dead *S. aureus* had no effect on HNO production of IL-1 family cytokines compared to uncolonized (**Fig. 5A**). Moreover, only live *S. aureus* increased apical IL-18 release and only live *S. aureus* increased basal HNO release of IL-1 family cytokines, with no detectable effect of live or dead *D. pigrum* or *S. pneumoniae*. In a human study of nasal inoculation with autologous *S. aureus*, increased IL-1β in nasal secretions was associated with *S. aureus* clearance from the nasal passage (IL-1α and IL-18 were not measured) (64). Thus, our data showing HNO release of IL-1 family cytokines in response to live *S. aureus* colonization recapitulates an *in vivo S. aureus* human nasal colonization response further supporting HNOs as a surrogate to identify host factors affecting *S. aureus* nasal colonization.

**Figure 5.**
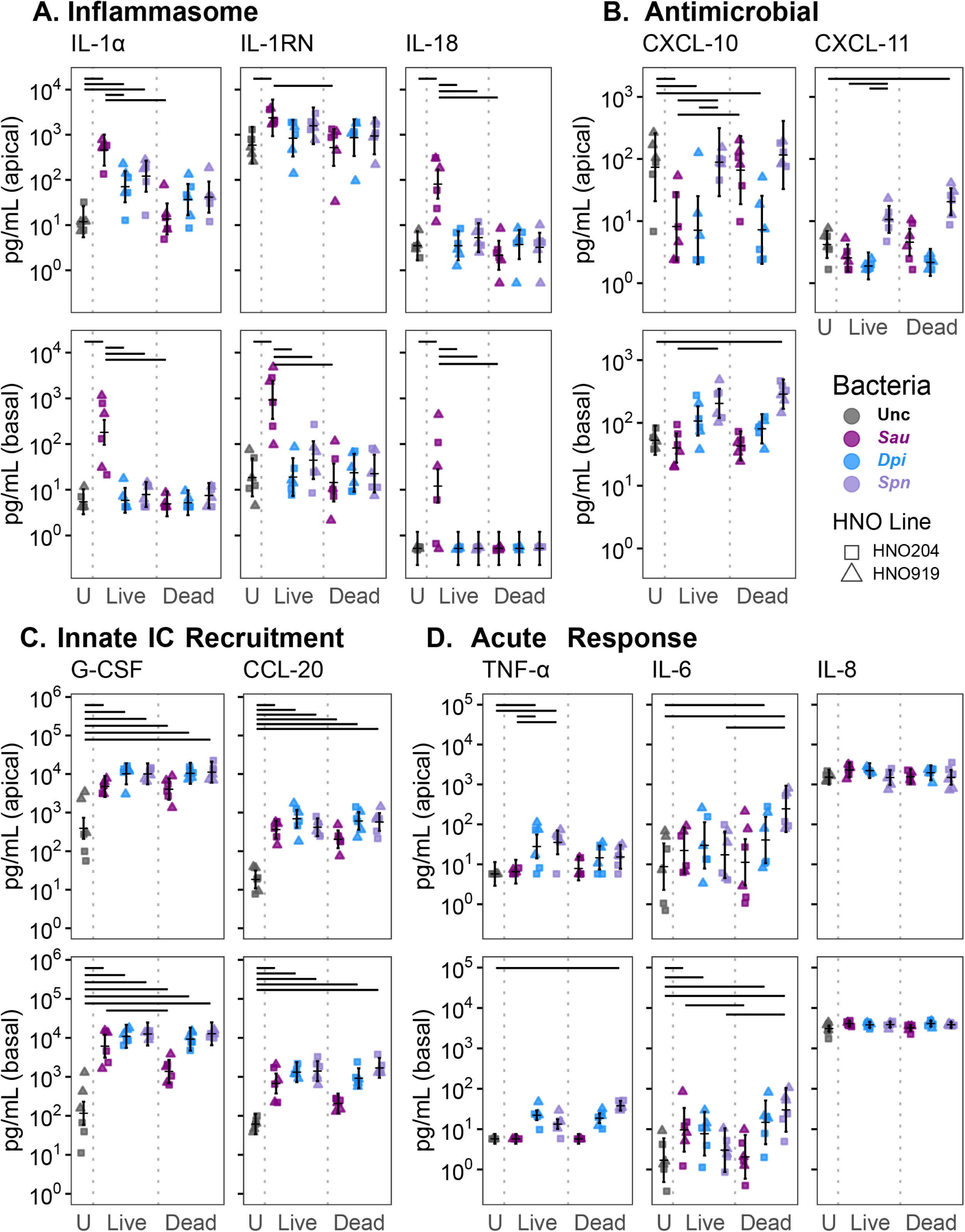
The human nasal respiratory epithelium of HNOs produces key cytokines in amounts orders of magnitude above the limit detection. HNO production of a subset of cytokines (pg/mL) detected in the apical wash or the basal medium after 48 h (shape indicates line and color indicates bacterial species). (**A**) Live *S. aureus* induced increased nasal epithelial production of IL-1 family cytokines (pg/mL) by 48 h, a response distinct from that to both *D. pigrum* and *S. pneumoniae*. (**B**) Live or dead *D. pigrum* and live *S. aureus* decreased epithelial apical production of CXCL10 compared to the uncolonized condition. In contrast, dead *S. pneumoniae* increased epithelial basal production of CXCL10 and apical production of CXCL11. (Basal CXCL11 was below limit of detection). (**C**) HNOs increased apical and basal production of G-CSF and CCL20 (MIP-3α) in response to all three bacteria, live or dead. (**D**) HNO production of TNF, IL-6, and IL-8 was modestly affected by bacterial monocolonization. Apical TNF mildly increased in response to live *D. pigrum* and *S. pneumoniae*, but not *S. aureus*. Apical IL-6 increased in response to dead *D. pigrum* or dead *S. pneumoniae.* Basal IL-6 increased in response to live *S. aureus*, live or dead *D. pigrum*, and dead *S. pneumoniae*. HNO production of IL-8 was stably high without and with bacterial colonization. Data are from three independent experiments in each of two HNO lines: HNO204 (squares), HNO919 (triangles). We used a LMM to determine statistical significance (**Table S1D-E**). The vertical brackets represent the model-predicted mean values and confidence intervals (+/- twice the standard error of the mean). Horizontal lines are included connecting pairs of bacterial conditions for all analyzed contrasts (see methods) that met the threshold for statistical significance (adjusted *p* < 0.05) and had a magnitude of effect greater than 4-fold (with this threshold chosen to highlight differences with a larger effect size and improve readability of the bars). See Table S1E for all adjusted *p* values.

Several monocolonization conditions resulted in decreased secretion of cytokines involved in antimicrobial immune responses below the levels of uncolonized controls (**Fig. 5B**). Live or dead *D. pigrum* or live *S. aureus* decreased apical production of CXCL10 (previously known as IP-10) ∼10-fold compared to uncolonized control HNOs, whereas there was little-to-no apical change in response to *S. pneumoniae* (**Fig. 5B**). CXCL10 binds to the CXCR3 receptor inducing many downstream functions, e.g., chemotaxis of CXCR3+ cells, regulation of cell growth and proliferation, and promotion of apoptosis (65). CXCL10 also exhibits antibacterial effects against USA300 *S. aureus* (66) and can stimulate release of *S. aureus* Protein A (67). CXCL10 promotes inflammation, which can have detrimental or beneficial effects on disease outcomes. For example, in adults, higher serum levels of CXCL10 are associated with increased severity of SARS-CoV2 infection (68) and higher levels of CXCL10 in nasal lavage fluid are associated with type 1 chronic rhinosinusitis (69). In contrast, in infants, higher levels of CXCL10 in nasal washes are associated with protection against severe respiratory syncytial virus infection (70). In young adults and children, nasal microbiota profiles enriched for *Dolosigranulum*/*Corynebacterium* are associated with lower likelihood of severe respiratory symptoms during SARS-CoV2 infections (71). It is interesting to speculate that *D. pigrum* could modulate CXCL10 levels during SARS-CoV2 infection since high levels of CXCL10 in SARS-CoV2 infection are associated with increased severity (68). Either live or dead *D. pigrum* suppressed nasal epithelial production of CXCL10, whereas only live, but not dead, *S. aureus* suppressed CXCL10 production (**Fig. 5B**), suggesting different mechanisms of suppression. Although inoculating peripheral blood mononuclear cells with *S. aureus* peptidoglycan lowers CXCL10 protein levels (72), the nasal epithelial response observed here required live *S. aureus*. In contrast, dead *S. pneumoniae* increased production of basal CXCL10 and apical CXCL11. Increased mucosal CXCL10 levels are associated with pneumococcal carriage in older adults (73). CXCL11 also acts via binding to the CXCR3 receptor, activating the same pathways as CXCL10 (74), and, along with CXCL10, is up-regulated in response to pneumococcal lung infection in mice (75). Overall, our data demonstrate that the HNOs are an excellent model for investigating nasal epithelial cytokine responses to bacterial colonization.

HNOs also displayed some general responses to bacterial colonization. For example, in response to live or dead bacteria, HNOs increased production of cytokines that interact with macrophages and neutrophils (**Fig. 5C**). Compared to uncolonized HNOs, both live and dead bacteria of all three species induced a 10-to-109-fold increase in apical and basal production of G-CSF and CCL20 (previously known as MIP-3α), except that dead *S. aureus* only induced a 3.4-fold increase in basal CCL20. CCL20 is a chemokine that attracts lymphocytes (76) and dendritic cells (77). G-CSF is a growth factor that modulates neutrophil activity and enhances their survival in the epithelium (78). In contrast to the increase in G-CSF and CCL20, bacterial colonization of HNOs induced only mild increases in the pleiotropic cytokines TNF and IL-6, e.g., live *D. pigrum* and *S. pneumoniae* induced a 5-to-6-fold increase in apical TNF (**Fig. 5D**). Although, dead *S. pneumoniae* induced a 28- and 18-fold increase in IL-6 apical and basal, respectively. These findings are in line with observations of *S. aureus* or *S. aureus* cell-free conditioned medium on pHNECs (79, 80) or bacterial colonization on immortalized cell lines (31, 32). In contrast to the low-level baseline production of TNF and IL-6, uncolonized HNOs produced high levels of IL-8 (ranging from 1,000 to 5,000 pg/mL) without an increase in response to bacterial colonization. Of note, monocolonized HNOs did not increase production of type 1 interferons (**Fig. 4**; **Table 1E**), a key epithelial response to bacterial and viral infection (81, 82), consistent with this being a model of microbiont colonization of the human nasal epithelium.

## DISCUSSION

To our knowledge, this is the first report using HNOs differentiated at an ALI to investigate colonization of nasal respiratory epithelium by common human nasal microbionts. HNOs differentiated at ALI (2D) have been used previously to investigate disease phenotypes, e.g., cystic fibrosis (83), including epithelial interactions with *Pseudomonas aeruginosa* (84), as well as respiratory viral infections (24–27). The differentiation at and maintenance of an ALI separates these HNOs from 3D HNOs and from 2D HNOs with apical medium (85, 86). This study also shows that colonizing bacteria restricted to the mucociliary blanket are sufficient to induce an innate immune response in nasal epithelium. Our results show that monocolonization of human nasal epithelium with three human nasal species resulted in distinct colonization dynamics and host responses. An epidemic USA300 strain of community-associated (CA) MRSA showed an initial drop in mucosal burden followed by a subsequent rebound indicating active replication on nasal epithelium. Moreover, the epithelial innate immune response to USA300 CA-MRSA included higher levels of IL-1 family cytokines, indicative of activation of an inflammasome response. We speculate that the physiological mucus layer of the HNOs might play a role in lengthening the time that HNOs tolerate colonization with USA300 CA-MRSA (clonal complex 8) compared to what is reported for Calu-3 cells polarized at ALI (31). Additionally, we demonstrated that the candidate nasal mutualist *D. pigrum* reduced human nasal epithelial production of CXCL10, an inflammatory cytokine implicated in antimicrobial response and immune activation. Nasal colonization by *D. pigrum* has repeatedly been associated with positive health outcomes and HNOs now provide a means to elucidate the mechanisms of *D. pigrum* interactions with human nasal epithelium. In contrast, *S. pneumoniae*, a nasal pathobiont associated with negative health outcomes and worsened viral infection symptoms, increased HNO production of antimicrobial and inflammation-associated cytokines. Unlike experimental human pneumococcal colonization, HNOs could be used to study colonization by a broader range and more virulent serotypes of *S. pneumoniae,* expanding efforts to identify key mechanisms of pneumococcal nasal colonization. Future applications for HNOs include extension to the pediatric nasal epithelium (25), microbe-epithelial interactions in specific diseases (e.g., CF), and the potential for genetic manipulation of organoid lines, as is done with human intestinal organoids (87). Of note, the FDA Modernization Act 2.0 permits the use of human organoids as preclinical models for function and toxicity based on the limitations of animals for modeling some human diseases and physiological characteristics (88). Overall, the data presented here demonstrate that HNOs are a powerful model system for elucidating microbiont-epithelial interactions and providing new insights into the functions of human nasal microbiota.

## MATERIALS and METHODS

### Isolation and passaging of 3D human nasal organoids (HNOs)

The isolation and passaging of human nasal epithelial organoids (aka human nose organoids) and the lines used here are previously described (24, 25). HNOs were established for use and distribution by the Baylor College of Medicine (BCM) 3D Organoid Core under a protocol (H-46014) approved by the BCM Institutional Review Board and with donor informed consent. In brief, both a nasal wash and a midturbinate swab were collected into a 50 mL conical tube from each donor and placed on ice until centrifugation. The supernatant was mixed with 10 mL airway organoid (AO) medium (as described in (24) without penicillin and streptomycin) containing 0.5 mg/ml collagenase (Sigma-C9407), 250 ng/mL amphotericin B, and 10 ng/mL gentamicin on an orbital shaker at 4 °C for 30 to 60 minutes (min). Three hundred μL of fetal bovine serum (FBS, Fisher #FB129999102) was added to the suspension to inactivate the collagenase and the suspension was mildly sheared using a 1 mL pipette tip and filtered through a 100-μm cell strainer (VWR #76327-102). The suspension was then centrifuged at 400 x g for 5 min at 4 °C (Eppendorf 5702R). Supernatant was discarded and the cell pellet was washed twice in 10 mL wash medium (as described in (24) without penicillin and streptomycin) and centrifuged at 400 x g for 5 min at 4 °C. Each nasal-cell pellet was resuspended in 30 μL of Corning Matrigel GFR Basement Membrane Matrix (Corning #356231), placed in a 24-well culture plate, and incubated at 37 °C, 5% CO_2_ for 20 min to solidify. Upon completed gelation, 500 μL of AO medium was added to each well and plates were incubated in a humidified 37 °C, 5% CO_2_ (Thermo Scientific Heracell Vios 160i). The medium was changed every 4 days. The cells were passaged every 7 days. To passage, the 3D HNOs were washed with 0.5 nM Ethylenediaminetetraacetic acid (EDTA, Thermo Scientific J15694AE) in cold Phosphate Buffered Saline (PBS) and centrifuged at 300 x g for 5 min at 4 °C. Each cell pellet was resuspended in 500 μL of 0.05% trypsin-EDTA (Gibco 25300120) then incubated at 37 °C in a 5% CO_2_ incubator for 4 min to break up the cells. To neutralize trypsin, we added 1 mL of wash medium with 10% FBS and dispersed the cells by pipetting 50 times. The cells were then centrifuged for 5 min at 300 x g and resuspended in 30 μL Matrigel per HNO to be generated. Cells were then placed into wells on a cell culture plate, allowed to solidify, and incubated as described above.

### Differentiation of HNOs as 2D cultures at an air-liquid interface

As previously published (24), after 7 days of propagation in AO medium, 3D HNOs were washed with 5 nM EDTA as above to disperse cells and then incubated for 5 min in 0.05% trypsin-EDTA. We then added 1 mL of wash medium with 10% FBS and dispersed the 3D HNOs into approximately single cells by mechanically pipetting 160 times with a P1000 tip. The single-cell suspension was filtered through a 40-μm VWR cell strainer (#76327-098), then centrifuged for 5 min at 400 x g at 4 °C. The cell pellet was resuspended in AO medium supplemented with 25 ng/mL Epidermal Growth Factor (EGF; Gibco #PMG8043) and 150 μL of the single-cell suspension was plated drop by drop on the center of the apical side of a cell-culture-coated transwell insert (Corning #3470) pretreated with 30 μg/mL of bovine Type I collagen (Gibco #A1064401). Then 600 μL of AO medium supplemented with 25 ng/mL EGF was added to the basal side of each transwell. (One 3D HNO well was used to seed one transwell.) HNOs were incubated at 37 °C in a humidified 5% CO_2_ incubator for 4 days, then medium was removed from both the basal and the apical sides and 600 μL of PneumaCult Airway Organoid Differentiation Medium (AODM; STEMCELL Technologies #05060) was added only to the basal side creating an air-liquid interface (ALI) culture. Basal medium was changed twice weekly while HNOs were differentiated at ALI for 21 days.

### RPMI 2650 cell cultivation at air-liquid interface

RPMI 2650 cells were purchased from ATCC (ATCC CCL-30) then grown and transitioned to ALI according to the method in Huffines et al. (33). Briefly, RPMI 2650 cells were cultivated in Minimal Essential Medium (MEM: Corning 10010CM) supplemented with 10% heat-inactivated (HI) FBS (Corning 35011CV), 1% L-glutamine (Gibco 25030081), and 1% Antibiotic-Antimycotic (AA: Gibco 15240062) in cell culture flasks in a 37 °C, 5% CO_2_, humidified incubator until 80-90% confluent. Transwell inserts were coated with 100 µL of Vitrogen plating medium (MEM without L-glutamine or phenol red (Gibco 51200038) supplemented with 10 µg/mL fibronectin (Advanced Biomatrix 5050-1MG), 100 µg/mL bovine serum albumin (VWR K719-50ML), and 30 µg/mL bovine collagen (Advanced Biomatrix 5006-15MG)) then placed in a UV crosslinker (Stratalinker 1800) for 45 mins prior to washing twice with 150 µL of sterile EBSS. Coated transwell inserts were seeded with 100 µL of RPMI 2650 cells at a concentration of 2.5 x 10^5^ cells/100 µL and 600 µL of culture medium was added to the basal side of the transwell then grown in a 37 °C, 5% CO_2_, humidified incubator for 7 days. The apical and basal medium were changed every other day. On the 7^th^ day of incubation, cells were transitioned to ALI by removing the apical medium and rinsing the apical side of the cells twice with 150 µL of EBSS. The basal side of the transwell was washed with antibiotic-free culture medium and transferred into 600 µL of antibiotic-free basal culture medium in a new sterile 24-well plate. Cells were incubated for 7 more days, with every other day medium changes. On day 14 after seeding, cells were transferred to a 34 °C, 5% CO_2_, humidified incubator for 24 hours (h) prior to experimentation.

### Calu-3 cell cultivation at air-liquid interface

Calu-3 cells were purchased from ATCC (ATCC HTB-55) then grown and transitioned to ALI according to the method in Kiedrowski et al. (31, 89). Cells were grown in base medium consisting of MEM supplemented with 10% HI FBS, 1% non-essential amino acid (NEAA) solution (Gibco 11140050), and 1% AA in cell culture flasks until 80-90% confluent. Prior to seeding, transwells were coated with 200 µL of filter-sterilized 60 µg/mL human placental collagen (Sigma C7521) for 24 h at room temperature. Then the collagen was removed and the transwells were dried in a biosafety cabinet for 15 mins, then washed twice by submerging each transwell in a 24-well plate well filled with 2 mL of EBSS (removing apical EBSS with a pipette). Confluent Calu-3 cells were seeded on transwells at 2.5 x 10^5^ cells/transwell in 100-to-200 µL of seeding medium consisting of DMEM:F-12 (Gibco 111320-033) supplemented with 5% HI FBS and 1% NEAA solution. Seeding medium (600 µL) was added to the basal side of the transwell. Cells were cultivated in a 37 °C, 5% CO_2_, humidified incubator for 24 h. At this point, the basal seeding medium was removed and replaced with polarization medium (DMEM:F12 supplemented with 0.5% HI FBS and 1% AA) and the cells were transitioned to ALI by removing the apical medium and rinsing the apical side of the cells once with 200 µL polarization medium before incubating in a 37 °C, 5% CO_2_, humidified incubator for 14 days. For the first 7 days, any liquid on the apical side was removed daily to maintain ALI. On day 7 (6 days at ALI), basal medium was replaced with antibiotic-free polarization medium, and the apical surface of the well was washed with 200 µL antibiotic-free polarization medium.

### Transepithelial electrical resistance (TEER)

The minimum TEER for each batch of HNOs used for experiments was 600 Ohms. The median (range) TEER for each line was as follows: HNO204 900 (655–1825) Ohms, HNO918 880 (585–1090) Ohms, HNO919 840 (700–1460) Ohms. We measured the TEER of 1 (and sometimes 2) individual HNO per line per experiment immediately after receiving wells from the Baylor College of Medicine 3D Organoid Core, which was ∼24 h before bacterial inoculation. We added 100 μL of Earle’s Balanced Salt Solution without calcium, magnesium, or phenol red (EBSS; Gibco 14155063) to the apical side of the selected HNO, placed the TEER meter probe (World Precision Instruments EVOM2 with STX2 electrodes) into the liquid in the transwell according to the methods in (90), and recorded the Ohms. After probe removal, cells were scraped off the transwell with a pipet tip, resuspended in 100 μL of EBSS, and transferred to a clean 1.5 mL microcentrifuge tube for mycoplasma testing. After testing TEER on an individual HNO well, we changed the basal medium of all the other HNO wells and placed the 24-transwell plate into a humidified 5% CO_2_ incubator at the specified temp (34 °C or 37 °C). The same methods were used for Calu-3 and RPMI 2650 cells with the exception that the TEER meter probe was a World Precision Instruments EVOM with STX4 electrodes. Calu-3 cells had a median TEER of 570 Ohms (range 500-650 Ohms) and RPMI 2650 cells had a median TEER of 350 Ohms (range 350-375).

### Mycoplasma testing

A representative well on each transwell plate for each HNO line was confirmed to be negative for *Mycoplasma* prior to experimentation as described here. The microfuge tube containing the resuspended HNO that had been used to measure TEER (above) was heated at 95 °C for 15 min to lyse HNO cells. From the same transwell, we collected the basal medium into a separate 1.5 mL microcentrifuge tube. Each tube was used in a separate PCR reaction with either the Biovision Mycoplasma PCR Detection kit or the Venor GeM Mycoplasma Detection kit, per manufacturer’s protocol. Electrophoresis on the PCR reactions on a 1% TopVision agarose gel made with and ran in a 1X Tris Acetate EDTA (TAE) buffer followed by staining DNA with SYBR Safe (Invitrogen #S33102) was used to detect which samples were positive for *Mycoplasma*. Only transwell plates with a representative HNO negative for *Mycoplasma* were used for experiments. These same steps were also performed for Calu-3 and RPMI 2650 cells.

### Epithelial RNA extraction and sequencing

HNOs were plated at ALI and differentiated at 37 °C for 21 days, as described above. On day 21, the 2D HNOs at ALI were either collected for RNA extraction (red in Fig. S1A) or shifted to 34 °C for two more days (grey in Fig. S1A). RNA was extracted from 2 pooled HNOs for each line from each condition. HNO wells were first washed once with 2x PBS. The following steps were then completed using the QIAGEN RNeasy Mini kit per the manufacturer’s instructions using the on-column DNase digestion and Qiashredder tubes. Briefly, we added 350 μL of RLT buffer with β-mercaptoethanol (BME; 10 μL BME per 1 mL RLT buffer) in the HNO well and mixed by pipetting then transferred the lysate into a QIAshredder column for centrifugation at 16000 x g for 2 min at room temperature (RT), discarding the QIAshredder column. We added 350 μL of freshly made 70% ethanol to the filtrate and mixed well by pipetting. The sample was then transferred to a RNeasy Mini spin column in a 2 mL collection tube and centrifuged at 16000 x g for 30 sec, discarding filtrate. After performing the On-Column DNase digestion step, we continued with RNA cleanup steps. To elute RNA, we placed the RNeasy spin column in a 1.5 mL collection tube, added 40 μL RNase-free water directly to the spin column membrane and centrifuged for 2 min at 16000 x g. A Nanodrop 2000 (Thermo Fisher Scientific) was used to quantify the RNA. A 260/230 ratio of ≥ 2 was the threshold for use for RNA sequencing. Novogene performed the sequencing using non-directional poly-A library preparation and 20 million paired reads using their Illumina NovaSeq 6000 and X-Plus Sequencing Platform.

### Analysis of RNAseq data

We used the FASTQC package v0.11.9 to inspect the quality of the raw sequence reads. We used TrimGalore v0.6.5 with the default settings to remove Illumina adapters and low-quality basepairs prior to aligning reads to human genome build GRCh38.98 using HiSAT2 v2.2.1 (91). We then used featureCounts to generate a count matrix from the aligned reads. The DeSeq2 package (v1.44.0) (92) was used to perform principal component analysis of the feature count data. Detailed code to analyze the RNAseq data and create the corresponding plot is available online (https://klemonlab.github.io/HNOBac_Manuscript).

### Enumeration of cells per transwell

The following method was used for HNOs, Calu-3 cells, and RPMI 2650 cells. To separate the epithelial cells from the transwell, we added 200 μL of 0.25% Trypsin-EDTA (Gibco #25200056) and incubated at 37°C in a humidified 5% CO_2_ incubator for 10-15 min. We then gently pipetted up and down 3 times with a 200 μL pipet tip to help dislodge the cells and incubated for an additional 10-15 minutes. We then gently pipetted up and down at least 20 times to disperse into individual cells and collected the suspension in a 1.5 mL Eppendorf tube. An aliquot of the epithelial cell suspension was mixed with equal parts Trypan blue. Viable HNO cells were counted manually using a Levy Double Count Chamber hemocytometer (Andwin Scientific 15170208) or using a BioRad TC20 automated cell counter (BioRad 1450102). The latter was used to count Calu-3 and RPMI 2650 cells. Detailed code to analyze the cells count data and create the corresponding plots is available online (https://klemonlab.github.io/HNOBac_Manuscript). Calu-3 cells had a median of 2.8 x 10^5^ cells/transwell and RPMI 2650 cells had a median of 4.5 x 10^6^ cells/transwell. HNO cells per transwell are shown in Figure S1B.

### Bacterial growth conditions

*Dolosigranulum pigrum* strain KPL3065 (93), the USA300 methicillin-resistant *Staphylococcus aureus* strain JE2 (94), and *Streptococcus pneumoniae* strain 603 (95) were each grown on BD BBL Columbia agar medium with 5% sheep’s blood (BAP) at 34°C in a humidified 5% CO_2_ incubator. We struck from a frozen 15% glycerol stock stored at −80 °C onto BAP for single colonies. After 36 h of growth, 10-15 *D. pigrum* single colonies were picked up with one sterile cotton swab (Puritan #25806) and bacteria were spread as a small lawn on one sterile 47 mm diameter, 0.2-μm polycarbonate membrane (Millipore Sigma #GTTP04700) atop BAP. This was repeated for four more BAP plates. After 36 h of growth, we collected *D. pigrum* from 3-4 membranes using a sterile cotton swab, then resuspended the cells in EBSS, and adjusted the optical density at 600 nm (OD_600)_ to yield ∼1×10^7^ CFUs per 15 μL (as described below). We struck MRSA JE2 from a frozen glycerol stock for single colonies on a sterile 47 mm, 0.2-μm polycarbonate membranes on BAP. After 14 h of growth, 5-10 colonies of MRSA were picked up with a sterile swab, resuspended cells in EBSS, and the resuspension was normalized to an OD_600_ that would yield ∼1×10^7^ CFUs per 15 μL EBSS. We struck *S. pneumoniae* from a frozen glycerol stock for singles directly onto BAP. After 12 h of growth, 10-15 colonies were picked up with a sterile cotton swab and swabbed as a lawn onto a sterile 47 mm, 0.2-micron polycarbonate membranes atop BAP, preparing 4 plates. After 14 h of growth, we collected *S. pneumoniae* lawns from 4 membranes on each side of a sterile swab, resuspended cells in EBSS, and adjusted the OD_600_ to yield ∼1×10^7^ CFUs per 15 μL EBSS.

### Calculation of the number of bacterial colony forming units in a given optical density

*D. pigrum* was grown for 36 h, and *S. aureus* and *S. pneumoniae* were grown for 12-14 h, as described above. For each species, 20 - 30 single colonies were resuspended in 2-3 mL of EBSS and the OD_600_ was measured. We then made a series of suspensions in EBSS with a range of OD_600_ values from 0.1 to 1.5 and performed serial dilutions of each of the suspensions. A drip-plate method was used to enumerate CFUs. We inoculated 20 μL from each dilution twice onto BAP, tilting the petri dish to allow the 20 μL to drip down the plate to facilitate the counting of colonies after growth. After counting colonies, we calculated the total CFUs per 15 μL of original suspension for each OD_600_ value and generated a standard curve to determine what OD_600_ would yield an HNO inoculum of 10^7^ CFUs/15 μL EBSS for each species.

### Bacterial colonization of epithelial cells and enumeration of bacterial CFUs

We also used the method described here for HNOs for both Calu-3 and RPMI 2650 cells. We gently pipetted 15 μL of each bacterial suspension (∼1×10^7^ CFUs) onto the surface of each corresponding HNO in a transwell. Uncolonized control HNOs received 15 μL of EBSS alone. The 24-well transwell plates were then centrifuged at 120 x g for 1 min at 25°C in an Eppendorf Centrifuge 5430 R to help absorb the bacterial suspension into the apical mucus layer. HNOs were then incubated at either 34°C or 37°C in a humidified 5% CO_2_ incubator. We collected HNOs for bacterial CFU enumeration after 24 or 48 h by first removing basal medium and adding 75 μL of 0.25% Trypsin-EDTA to the apical surface of each well and incubating at 37 °C for 15 min to lift the HNO from the transwell. Second, we added 75 μL of 0.025% Triton-X 100 (Thermo Scientific J66624.AE) to each well and then pipetted the mixture up and down 30 times to fully break up the epithelial cell layer. We then made serial dilutions to enumerate the total bacterial CFUs on each HNO. In a small subset of the independent experiments, *S. aureus* destroyed the HNO epithelial cell layer and the data from such wells was excluded from further analysis. Detailed code to analyze the CFU data, perform statistical analysis, and create the corresponding plots is available online (https://klemonlab.github.io/HNOBac_Manuscript).

### Lactate Dehydrogenase (LDH) Assay

We also used the method described here for HNOs for both Calu-3 and RPMI 2650 cells. Basal medium was collected at −1 h and at either 6 or 48 h relative to time of colonization from all HNO wells, including uncolonized controls, and immediately stored at −80 °C. LDH activity was measured in duplicate using the Promega CytoTox Non-Radioactive Cytotoxicity Assay (#G1780), per the manufacturer’s instructions. Cytotoxicity for each experiment was calculated as the fold change in LDH in basal medium from an HNO at either 6 or 48 h compared to the same HNO at −1 h relative to colonization, including for the uncolonized controls. Detailed code to analyze the LDH data, perform statistical analysis, and create the corresponding plots is available online (https://klemonlab.github.io/HNOBac_Manuscript).

### Brightfield Microscopy

Uncolonized HNOs, Calu-3 cells, or RPMI 2650 cells were inoculated with 15 μL of EBSS then incubated at 34 °C for 24 h (Fig. 1A) or 6 h (Fig. 2). The method described here for HNOs was also used for both Calu-3 and RPMI 2650 cells. We fixed HNOs in Clarke’s solution (75% ethanol and 25% glacial acetic acid) by removing each HNO-containing transwell from its plate and submerging it in 2 mL of RT fixative in the well of a 24-well plate for 30 min. After fixation, HNO transwells were rinsed twice by submersion in 100% anhydrous methanol at 4 °C for ≥ 30 min. We then cut the bottom membrane, on which the HNO sits, of each transwell from its plastic housing with a sterile scalpel and stored the HNO plus membrane at 4 °C in anhydrous methanol until submission to the TMC Digestive Disease Center Tissue Analysis & Molecular Imaging Core for paraffin embedding, sectioning, and staining. Sections were stained with Sigma-Aldrich Microscopy PAS staining kit (#1.01646.0001), and then counterstained with hematoxylin (Epredia, #7211). Images were taken with a Nikon Eclipse Ci-L bright field microscope (Serial Number 702085) at 40x magnification. Fiji Is Just ImageJ (FIJI) was used to enhance contrast, white balance the image, and add a scale bar.

### Fluorescence Microscopy

HNOs were inoculated with ∼1×10^7^ CFUs of *D. pigrum*, *S. aureus*, or *S. pneumoniae* in 15 μl of EBSS or, as a control, with EBSS alone, as described above. Live bacterial resuspensions were prestained with MitoTracker Red CMXRos (Invitrogen #M7512) before colonization (49). Briefly, each bacterium was resuspended in 10-15 mL of EBSS to an OD_600_ of 1 prior to adding 1 μL of 1 mM MitoTracker Red CMXRos per mL of resuspension and incubating at RT for 1 h. Prestained bacteria were washed twice with 10-15 mL of EBSS by centrifugation at 10000 x g for 10 min at RT and resuspended in EBSS. The OD_600_ of each resuspension was adjusted to yield ∼1×10^7^ CFUs per 15 μL of EBSS and HNOs were monocolonized with bacteria as described above. After 6 h of colonization at 34 °C, we fixed HNOs, including uncolonized controls, in Clarke’s solution, and stored them as described above. We used the same methods for uncolonized Calu-3 and RMPI 2650 cells as for uncolonized HNOs. Samples were submitted to the TMC Digestive Disease Center Tissue Analysis & Molecular Imaging Core for paraffin embedding, sectioning, and staining. Briefly, after tissue processing, the transwell membranes were bisected and embedded in paraffin (Richard-Allen Scientific, 22900700) for cross-sectioning. The paraffin-embedded membranes were deparaffinized and subjected to antigen retrieval (Biocare, CB910M). The sections were incubated with 4% bovine serum for 1 h to block nonspecific protein binding, then were incubated overnight at 4°C with primary antibodies to MUC5AC (Invitrogen, #MA5-12178, dilution 1:1000), KRT5 (Biolegend, #905503, dilution 1:1500), and/or acetylated alpha tubulin (Santa Cruz (6-11B-1), #sc-23950, dilution 1:1000). The primary antibodies for MUC5AC and acetylated alpha tubulin were detected using an Alexa Fluor 488-conjugated goat anti-mouse secondary antibody (Invitrogen #A11001; dilution 1:300) and for KRT5 was detected using an Alexa Fluor 568 goat anti-rabbit secondary antibody (Invitrogen # A11011; dilution 1:300). All slides were counterstained with Hoechst (Invitrogen™ 33258 #H3569, dilution 1:10,000) for 10 min and washed with deionized (DI) water. Coverslips were added with Invitrogen ProLong Glass Antifade Mountant (#P36980). Stained epithelial cells were imaged using an Olympus IX83 epifluorescence deconvolution microscope (#2000 IXplore IX83) equipped with an IX3 laser (#IX3-ZDC2) at 60x magnification using the blue (excitation 350 +/- 25 nm, emission 460 +/- 25 nm) and green (excitation 470 +/- 20 nm, emission 525 +/- 25 nm) channels to detect the Hoechst and MUC5AC signal, respectively (Fig. 1C-F). Stained HNOs were imaged at 100x magnification using the blue, green, and red (excitation 560 +/- 20 nm, emission 630 +/- 37.5 nm) channels to detect the Hoechst, MUC5AC, and MitoTracker Red CMXRos signal, respectively (Fig. 1C-F). For Figure 2 (D-I), uncolonized HNOs, Calu-3 cells, and RPMI 2650 cells were imaged at 60x magnification using the blue, red, and green channels to detect the Hoechst, KRT5, and alpha-tubulin/MUC5AC signal, respectively. All images were processed for publication using FIJI to adjust contrast to maximize visibility and minimize overexposure in each color channel separately, merge channels, and add scale bars.

For Figure 2, all microscopy images of HNOs, Calu-3 cells, and RPMI 2650 cells were processed identically to enable direct comparison of MUC5AC, KRT5, and alpha-tubulin staining. Images were acquired using an Olympus IX83 epifluorescence deconvolution microscope with consistent acquisition settings across all samples and channels. We set the minimum exposure time for each channel based on the HNO samples, and all images acquired from Calu-3 and RPMI 2650 cells used exposure times greater than or equal to that of the HNO samples. Maximum intensity projection images were generated from deconvolved z-stacks and used for analysis. Image processing was performed in FIJI. Background signal was subtracted using the built-in “Subtract Background” function with a rolling ball radius of 50 pixels. For visualization purposes, channel-specific contrast enhancement was applied using the “Enhance Contrast” function. For the Hoechst and MUC5AC channels, contrast was adjusted to 0.4% saturated pixels. For the KRT5 and alpha-tubulin channels, contrast was adjusted to 0.05% saturated pixels. All contrast adjustments were applied uniformly across comparable images to ensure consistency.

### Gamma-irradiation to kill bacteria

To sufficiently inactivate bacteria while maintaining structural integrity, we irradiated each bacterium with between > 1,000 and < 8,000 grays, per Correa et al. (59). We resuspended each bacterial species separately in EBSS to an OD_600_ equal to ∼1×10^7^ CFUs/15 μL and added 25 mL of each bacterial resuspension separately to a 50 mL conical tube and irradiated overnight in a Gammacell-1000 Irradiator (Atomic Energy of Canada Ltd.) with a Cesium-137 source. This instrument generates 758.3 rads/min, and each sample was irradiated for ∼9.5 h for a total of 4,322 grays of radiation. After irradiation, we gently washed bacteria to remove lingering reactive oxygen species by pelleting cells at 6,000 x g for 10 min, removing the supernatant, and resuspending in an equal amount of EBSS. We plated 2 drips of 20 μL of each resuspension on BAP and cultured for 48 h at 34°C in a humidified 5% CO_2_ incubator and verified the absence of viable bacteria. We aliquoted the remaining resuspension in 1 ml aliquots and flash-froze each in a dry ice and ethanol mix before storage at −80 °C. For each experiment, we thawed a fresh aliquot of gamma-irradiated bacteria at RT, and we used 15 μL of each resuspension to inoculate an HNO with dead bacteria. Two 20 μL drips were plated at the beginning of each experiment to verify the absence of viable bacteria in the experimental inoculum.

### Immunoassays for detection of cytokines

HNOs were colonized with either live (approximately 10^7^ CFUs) or dead (gamma-irradiated) bacteria resuspended in EBSS and included uncolonized control HNOs inoculated with buffer alone. After 48 h at 34°C in a humidified CO_2_ incubator, basal medium was collected from each well and transferred to a sterile 1.5 mL microcentrifuge tube. The apical side of each HNO was separately washed by adding 150 µL of AODM and pipetting up and down 3 times before transferring the wash to a sterile 1.5 mL microcentrifuge tube. Each HNO was washed a second time with the same volume and both washes were pooled. Samples were frozen at −80 °C until use in immunoassays. For immunoassays, samples were submitted to the Digestive Disease Center Functional Genomics and Microbiome core with the following Millipore Sigma magnetic-bead panel kits: 1) Milliplex Human Cytokine Panel III with IL-29, I-TAC, CCL20/MIP-3a, MIP-3b; 2) Milliplex Human Cytokine Panel A with MIG, IFNg, IFNa2, TNF-a, IP-10, RANTES, IL-1a, IL-1ra, IL-1b, IL-5, IL-6, IL-8, IL-10, IL-12p70, IL-13, IL-18, MIP-1a, MIP-1b, MCP-1, MCP-3, IL-17A, IL-17E/IL-25, Eotaxin/CCL11, G-CSF, GM-CSF, VEGF-A; 3) Milliplex Human MMP Panel 2 with MMP1, MMP2, MMP7, MMP9 and MMP10; and 4) Milliplex Human Cytokine Panel IV with IL-32, IL-36/IL-1F8, IL-37/IL-1F7, IL-38/IL-1F10. Samples were assayed per the manufacturer’s instructions and analyzed with Luminex xPONENT for Magpix (version 4.2, build 1324) on a Magpix instrument. Data was analyzed with Milliplex Analyst (version 5.1.0.0, standard build 10/27/2012). All sample values below the manufacturer’s limit of detection were set at the limit of detection for analysis. The set of assay results for a given cytokine at a given location (e.g., apical IL-6) across all inoculation conditions was considered below the limit of detection (gray tiles in Fig. 4) and excluded from subsequent statistical analysis if > 75% of those results were at or below the manufacturer’s limit of detection, except when the ≤ 25% of those samples that were above that threshold had log_2_-fold changes for colonized compared to uncolonized that were ≥ 2.5 times higher than the limit of detection. This only occurred for basal production of IL-18 in response to live MRSA, since no other condition stimulated much IL-18 release basally. Detailed code to analyze the cytokine data, perform statistical analysis, and create the corresponding plots is available online (https://klemonlab.github.io/HNOBac_Manuscript).

### Statistical Analysis

R (v4.4.0) and RStudio (v2023.12.1+402) were used for data analysis, statistics, and data visualization (96, 97). A complete list of the R packages used is available at https://klemonlab.github.io/HNOBac_Manuscript/RSession.html. For analysis of the CFU, LDH, and cytokine data, we used the lme4 package (v1.1-35.5) to fit the data to linear mixed-effects models (98). For the CFU and LDH data, we used the Holm method to adjust for multiple comparisons (99). For each assayed HNO temperature, we modeled the log-transformed CFU counts considering bacterial species and collection timepoint as fixed effects and comparing results within each bacterial species between the 0-to-24 h and 24-to-48 h timepoints (**Table S1A**). When comparing CFU results between both temperatures, we considered as fixed effects both temperature and an interaction variable describing bacterial condition at each time point and analyzed only for contrasts across temperatures (**Table S1A**). When comparing CFU results between epithelial models, we considered bacterial condition and epithelial model as fixed effects and performed pairwise comparisons between the different epithelial models for each bacterium (**Table S1A**). Likewise, at each HNO temperature, we modeled the ratio of LDH activity after 48 h of colonization relative to 1 h before colonization with bacterial species as a fixed effect and, then, compared each species to the uncolonized condition (**Table S1B**). When comparing LDH results between both temperatures, we considered as fixed effects both temperature and bacterial condition and analyzed only contrasts across the temperatures (**Table S1B**). When comparing LDH results between epithelial models, we considered bacterial condition and epithelial model as fixed effects and performed pairwise comparisons between the different epithelial models for each bacterium and in the uncolonized condition (**Table S1B**). We modeled the log-transformed pg/mL data for each different cytokine assay derived from the apical or basal surfaces of the HNOs separately defining as a fixed effect in each model an interaction variable that combines bacterial condition (uncolonized, *D. pigrum, S. aureus*, or *S. pneumoniae*) with viability status (control, live, or dead). Across all the analyzed data we considered experimental date (i.e., independent experiment) nested under HNO line as random effects (**Tables S1A, S1B, and S1D**). This identifies and corrects for the percentage of variation due to donor line differences and experimental batch effects. Some of the cytokine’s fitted models were ‘singular’ due to HNO line-associated variance being close to zero; those models were recalculated with only experimental date as a random effect. We performed an ANOVA for each model with *p*-values added by the afex package (v1.4-1) (100). If the resultant global *p*-value for fixed effects was < 0.05, then individual contrasts were calculated with the emmeans package (v1.10.4) (101). For the cytokine data, we analyzed the following contrasts: the uncolonized control and all other conditions, all combinations of live bacterial species, and live vs. dead for each bacterium. To account for the large number of comparisons (588 contrasts), we then used false discovery rate (FDR) (102) for *p*-value adjustment across all the selected contrasts across all cytokine assays (**Table S1E**). Detailed code for all statistical analysis is available https://klemonlab.github.io/HNOBac_Manuscript.

## Data Availability

Sequencing data files and raw count matrix file are available at the Gene Expression Omnibus, accession number GSE277582. The data frames and code for all of the figures are available at https://klemonlab.github.io/HNOBac_Manuscript. The rest of the microscopy images for Figures 1 and 2 are available at https://github.com/KLemonLab/HNOBac_Manuscript/tree/main/data/microscopy.

## Acknowledgements

We thank the donors who make HNOs possible. We thank Sue Crawford for performing gamma irradiation of bacteria; Victoria Poplaski for RNA extraction of uncolonized HNOs; Scott Chimileski for microscopy advice; Pamela Parsons for microscopy staining; Elina Mosa for assistance with microscopy imaging; Robert Britton, Anthony Maresso, members of the Respiratory Organoid Group and of the Lemon Lab for helpful discussions. This research was supported by funding from the National Institute of Allergy and Infectious Diseases of the National Institutes for Health (NIH) under award U19AI157981 (K.P.L., K.A.P., S.E.B.), U19AI144297 (S.E.B), U19AI116497 (P.A.P., S.E.B.), and F31AI172324 (A.I.B.). Services from the Texas Medical Center Digestive Disease Center Functional Genomics and Microbiome Core (FGM) and Tissue Analysis & Molecular Imaging Core (TAMI) were supported in part by NIH grant P30DK056338. Biostatistics services from the Baylor College of Medicine (BCM) Biostatistics and Informatics Shared Advanced Technology Core were underwritten by institutional funds from BCM. The BCM Integrated Microscopy Core is supported by the Center for Advanced Microscopy and Image Informatics (CAMII) with funding from NIH (DK56338, CA125123, ES030285), and CPRIT (RP150578, RP170719).

## Author Contributions

Conceptualization: A.I.B., L.A.K., K.P.L. Methodology: A.I.B., L.A.K., I.F.E., H.T., S.G.H., A.K., A.R., K.A.P., P.A.P., S.E.B., J.M.L. Investigation: L.A.K., A.I.B., A.K. Resources: A.K., S.E.B. Data curation. I.F.E. Formal analysis. I.F.E., A.I.B., L.A.K., H.N.P., S.G.H. Validation. L.A.K., A.I.B., J.M.L. Visualization: A.I.B., L.A.K., I.F.E., S.G.H., H.N.P Writing – original draft: A.I.B., L.A.K., I.F.E., K.P.L. Writing – review & editing: L.A.K., A.I.B., I.F.E., K.P.L., K.A.P., S.G.H., A.K., H.T., H.N.P., A.R., J.M.L., P.A.P., S.E.B. Funding acquisition: K.P.L., S.E.B., A.I.B. Supervision: I.F.E., K.P.L.

## Supplemental Information

**Figure S1.**
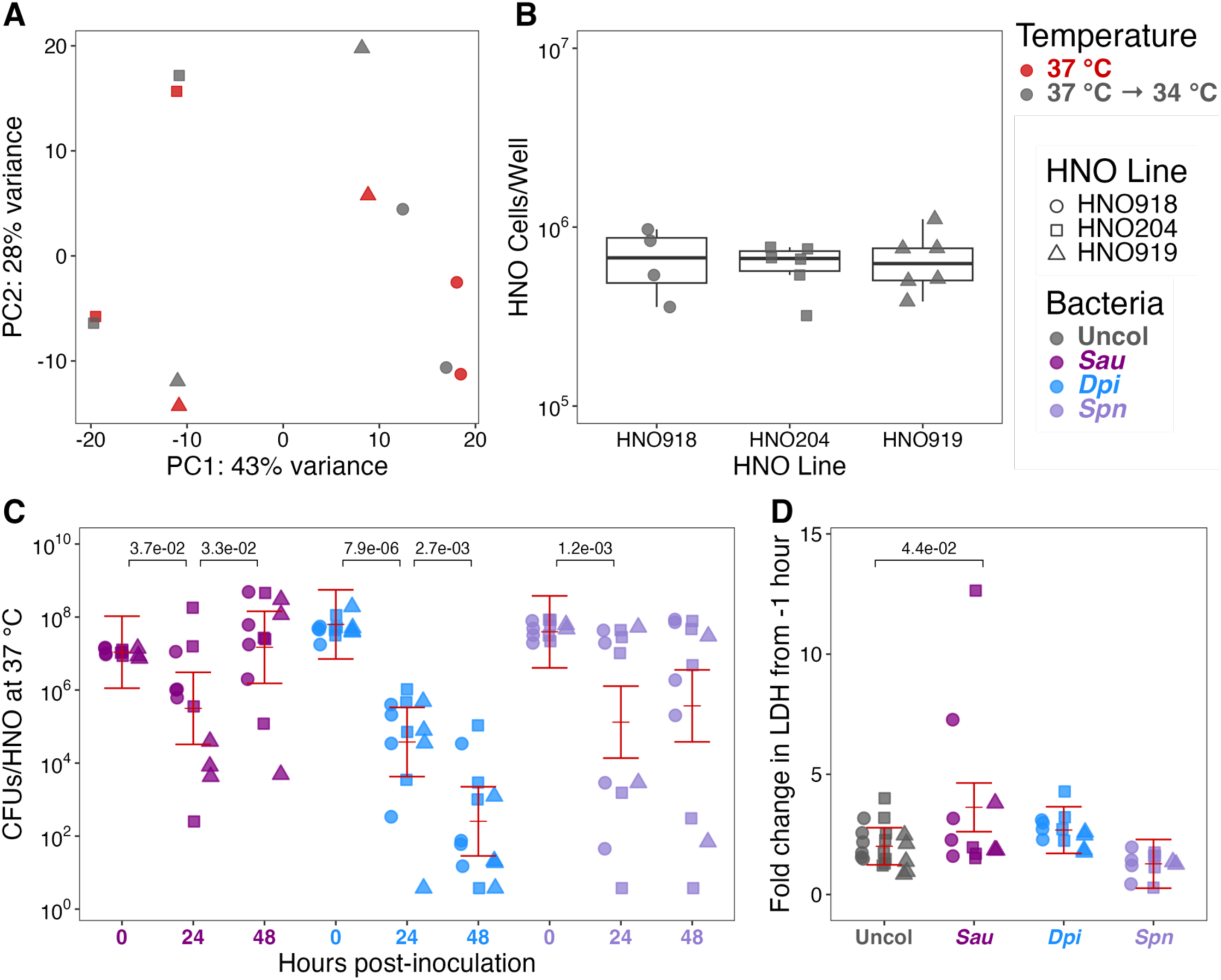
Nasal microbionts colonize HNOs at human internal body temperature, 37 °C. (**A**) In a Principal Component Analysis, epithelial transcription (read counts) within an HNO line was comparable between HNOs differentiated at 37 °C for 21 days (red) and HNOs shifted to 34 °C (gray) for an additional 2 days after the 21 days of differentiation at 37 °C (*n* = 2 independent experiments with 3 HNO lines). (**B**) HNO lines derived from different donors had comparable cells per transwell. The median (range) number of cells in each HNO was 6.9 x 10^5^ for line HNO918 (3.58 x 10^5^ – 9.72 x 10^5^), 6.68 x 10^5^ (3.2 x 10^5^ – 7.72 x 10^5^) for line HNO204, and 6.37 x 10^5^ for line HNO919 (3.84 x 10^5^ – 1.1 x 10^6^) in *n* = 4 for HNO918, *n* = 6 for HNO204, and *n* = 6 in HNO919. (**C**) HNOs were monocolonized with *S. aureus* (purple), *D. pigrum* (blue), and *S. pneumoniae* (lavender) at 37 °C for up to 48 h. At time 0, 10^7^ CFUs of a bacterium in 15 µL of EBSS were inoculated apically. Recovered CFUs/HNO at 24 and 48 h are shown. (**D**) Fold change in lactate dehydrogenase release (LDH) of uncolonized (gray) HNOs and of HNOs colonized with *S. aureus* (purple), *D. pigrum* (blue), or *S. pneumoniae* (lavender) into HNO basal medium at 48 h compared to −1 h samples from the same well at 37 °C. HNOs colonized by *S. aureus* had 1.8-fold higher basal LDH release compared to the uncolonized control. For **C** and **D,** the independent experiments per HNO line were HNO918 ≥ 4, HNO204 ≥ 4, and HNO919 ≥ 2. Data (**C, D**) were analyzed using a LMM (**Tables S1A-B)** to determine statistical significance and the Holm method was used to adjust *p*-values (shown above the horizontal bars) for multiple comparisons (0 to 24 h and 24 to 48 h in C and uncolonized to each bacterial treatment in D). Vertical brackets represent the model-predicted mean values and confidence intervals (+/- twice the standard error of the mean).

**Figure S2.**
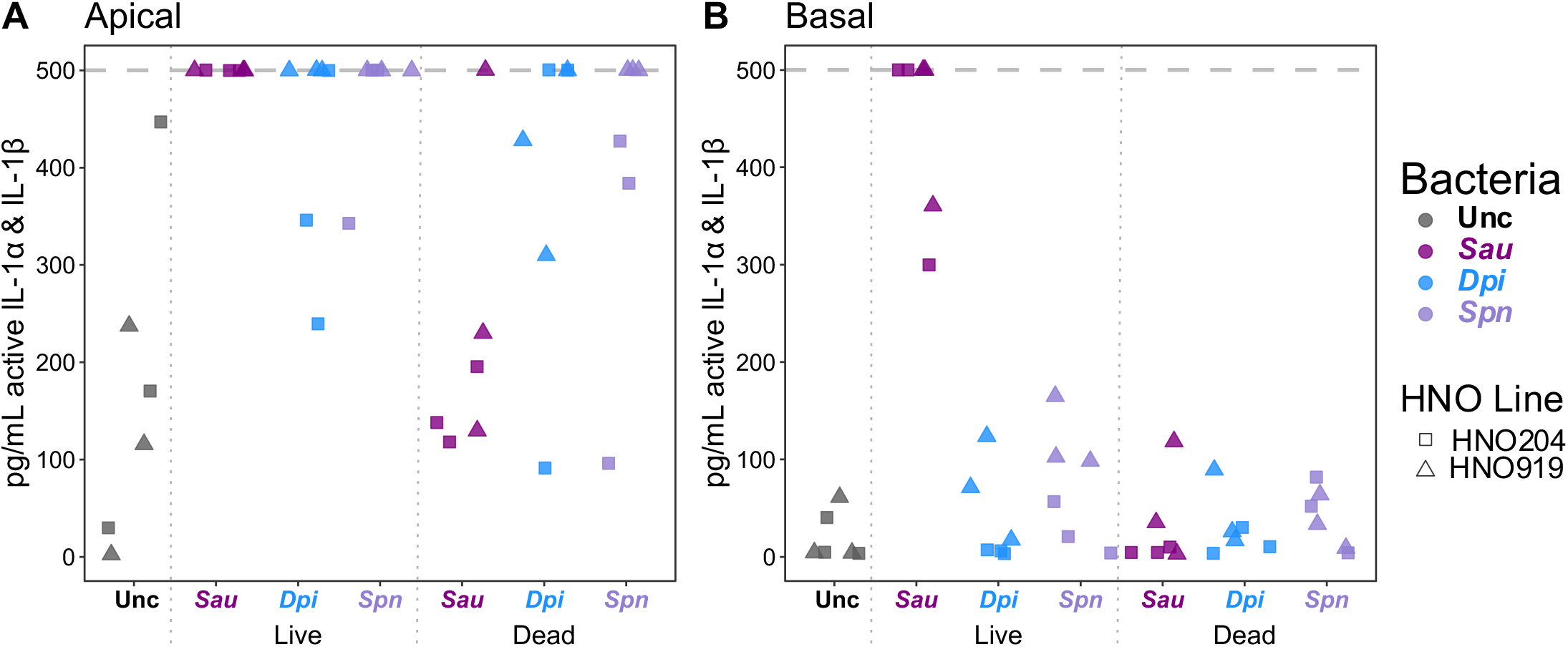
In response to live *S. aureus* monocolonization, HNO production of IL-1α activity dominates over IL-1RN activity resulting in IL-1 receptor activation apically and basally. (**A**) The apical and (**B**) basal IL-1α and IL-1β detected in the cytokine assays in Figure 5A are active. HEK-Blue™ IL-1R Cells were used to detect catalytically active IL-1α and IL-1β produced by HNOs (without distinction) to the extent that these were greater than IL-1RN activity. The upper limit of detection of the assay was 500 pg/mL (gray dashed line). For A, B, and C, data are from three independent experiments in each of two HNO lines (HNO204, HNO919). HEK-Blue™ IL-1R Cells (InvivoGen HKB-IL1R) were propagated and assays were performed according to the manufacturer’s instructions. Briefly, after adding 20 µL of either HNO basal medium or apical wash to the bottom of a 96-well plate, we inoculated each well with 3 x 10^5^ HEK-Blue cells and incubated for 22 hours at 37 °C in a humidified 5% CO_2_ incubator. HEK-Blue cells partially adhere to the bottom of the 96-well plate during incubation. At collection, plates were tapped gently to mix the liquid and 20 µL of spent HEK medium from each well was transferred into a corresponding well of a new 96-well plate prior to addition of 180 μL of freshly prepared QUANTI-Blue™ Solution (Invivogen REPQBS) per well. The assay developed at 37 °C for 2.5 hours before reading absorbance at 620 nm. A standard curve was constructed using InvivoGen recombinant human IL-1β as a positive control. Detailed code to analyze the HEK-Blue data and create the plots is available online (https://klemonlab.github.io/HNOBac_Manuscript/Methods_HekBlue.html).

Table S1. Results from linear mixed-effect models

## REFERENCES

1. Antimicrobial Resistance C. 2022. Global burden of bacterial antimicrobial resistance in 2019: a systematic analysis. Lancet 399:629–655. 10.1016/S0140-6736(21)02724-0.

2. Kluytmans J, van Belkum A, Verbrugh H. 1997. Nasal carriage of *Staphylococcus aureus*: epidemiology, underlying mechanisms, and associated risks. Clinical microbiology reviews 10:505–520.

3. Gorwitz RJ, Kruszon-Moran D, McAllister SK, McQuillan G, McDougal LK, Fosheim GE, Jensen BJ, Killgore G, Tenover FC, Kuehnert MJ. 2008. Changes in the prevalence of nasal colonization with *Staphylococcus aureus* in the United States, 2001-2004. J Infect Dis 197:1226–1234. 10.1086/533494.

4. von Eiff C, Becker K, Machka K, Stammer H, Peters G, Group FtS. 2001. Nasal carriage as a source of *Staphylococcus aureus* bacteremia. N Engl J Med 344:11–16.

5. Wertheim HF, Vos MC, Ott A, van Belkum A, Voss A, Kluytmans JA, van Keulen PH, Vandenbroucke-Grauls CM, Meester MH, Verbrugh HA. 2004. Risk and outcome of nosocomial *Staphylococcus aureus* bacteraemia in nasal carriers versus non-carriers. Lancet 364:703–705. 10.1016/S0140-6736(04)16897-9.

6. Young BC, Wu CH, Gordon NC, Cole K, Price JR, Liu E, Sheppard AE, Perera S, Charlesworth J, Golubchik T, Iqbal Z, Bowden R, Massey RC, Paul J, Crook DW, Peto TE, Walker AS, Llewelyn MJ, Wyllie DH, Wilson DJ. 2017. Severe infections emerge from commensal bacteria by adaptive evolution. Elife 6. 10.7554/eLife.30637.

7. Missiakas D, Schneewind O. 2016. Staphylococcus aureus vaccines: Deviating from the carol. J Exp Med 213:1645–1653. 10.1084/jem.20160569.

8. Miller LS, Fowler VG, Shukla SK, Rose WE, Proctor RA. 2020. Development of a vaccine against Staphylococcus aureus invasive infections: Evidence based on human immunity, genetics and bacterial evasion mechanisms. FEMS Microbiol Rev 44:123–153. 10.1093/femsre/fuz030.

9. Bode LG, Kluytmans JA, Wertheim HF, Bogaers D, Vandenbroucke-Grauls CM, Roosendaal R, Troelstra A, Box AT, Voss A, van der Tweel I, van Belkum A, Verbrugh HA, Vos MC. 2010. Preventing surgical-site infections in nasal carriers of *Staphylococcus aureus*. N Engl J Med 362:9–17. 10.1056/NEJMoa0808939.

10. van Rijen M, Bonten M, Wenzel R, Kluytmans J. 2008. Mupirocin ointment for preventing *Staphylococcus aureus* infections in nasal carriers. Cochrane Database Syst Rev:CD006216. 10.1002/14651858.CD006216.pub2.

11. Nair R, Perencevich EN, Blevins AE, Goto M, Nelson RE, Schweizer ML. 2016. Clinical Effectiveness of Mupirocin for Preventing Staphylococcus aureus Infections in Nonsurgical Settings: A Meta-analysis. Clin Infect Dis 62:618–630. 10.1093/cid/civ901.

12. Richardson EJ, Bacigalupe R, Harrison EM, Weinert LA, Lycett S, Vrieling M, Robb K, Hoskisson PA, Holden MTG, Feil EJ, Paterson GK, Tong SYC, Shittu A, van Wamel W, Aanensen DM, Parkhill J, Peacock SJ, Corander J, Holmes M, Fitzgerald JR. 2018. Gene exchange drives the ecological success of a multi-host bacterial pathogen. Nat Ecol Evol 2:1468–1478. 10.1038/s41559-018-0617-0.

13. Simell B, Auranen K, Kayhty H, Goldblatt D, Dagan R, O’Brien KL, Pneumococcal Carriage G. 2012. The fundamental link between pneumococcal carriage and disease. Expert Rev Vaccines 11:841–855. 10.1586/erv.12.53.

14. CDC. 2019. Antibiotic Resistance Threats in the United States, 2019. U.S. Department of Health and Human Services, CDC, Atlanta, GA.

15. Robinson RE, Myerscough C, He N, Hill H, Shepherd WA, Gonzalez-Dias P, Liatsikos K, Latham S, Fyles F, Doherty K, Hazenberg P, Shiham F, McLenghan D, Adler H, Randles V, Zaidi S, Hyder-Wright A, Mitsi E, Burhan H, Morton B, Rylance J, Lesosky M, Gordon SB, Collins AM, Ferreira DM. 2023. Comprehensive review of safety in Experimental Human Pneumococcal Challenge. PLoS One 18:e0284399. 10.1371/journal.pone.0284399.

16. Stubbendieck RM, Hurst JH, Kelly MS. 2024. *Dolosigranulum pigrum*: A promising nasal probiotic candidate. PLoS Pathog 20:e1011955. 10.1371/journal.ppat.1011955.

17. Escapa IF, Chen T, Huang Y, Gajare P, Dewhirst FE, Lemon KP. 2018. New Insights into Human Nostril Microbiome from the Expanded Human Oral Microbiome Database (eHOMD): a Resource for the Microbiome of the Human Aerodigestive Tract. mSystems 3:e00187–00118. 10.1128/mSystems.00187-18.

18. Brugger SD, Eslami SM, Pettigrew MM, Escapa IF, Henke MT, Kong Y, Lemon KP. 2020. *Dolosigranulum pigrum* Cooperation and Competition in Human Nasal Microbiota. mSphere 5:e00852-00820. 10.1128/mSphere.00852-20.

19. Liu CM, Price LB, Hungate BA, Abraham AG, Larsen LA, Christensen K, Stegger M, Skov R, Andersen PS. 2015. *Staphylococcus aureus* and the ecology of the nasal microbiome. Sci Adv 1:e1400216. 10.1126/sciadv.1400216.

20. Laufer AS, Metlay JP, Gent JF, Fennie KP, Kong Y, Pettigrew MM. 2011. Microbial communities of the upper respiratory tract and otitis media in children. mBio 2:e00245–00210. 10.1128/mBio.00245-10.

21. Bomar L, Brugger SD, Yost BH, Davies SS, Lemon KP. 2016. *Corynebacterium accolens* Releases Antipneumococcal Free Fatty Acids from Human Nostril and Skin Surface Triacylglycerols. mBio 7:e01725–01715. 10.1128/mBio.01725-15.

22. De Boeck I, Wittouck S, Martens K, Spacova I, Cauwenberghs E, Allonsius CN, Jorissen J, Wuyts S, Van Beeck W, Dillen J, Bron PA, Steelant B, Hellings PW, Vanderveken OM, Lebeer S. 2021. The nasal mutualist *Dolosigranulum pigrum* AMBR11 supports homeostasis via multiple mechanisms. iScience 24:102978. 10.1016/j.isci.2021.102978.

23. Sachs N, Papaspyropoulos A, Zomer-van Ommen DD, Heo I, Bottinger L, Klay D, Weeber F, Huelsz-Prince G, Iakobachvili N, Amatngalim GD, de Ligt J, van Hoeck A, Proost N, Viveen MC, Lyubimova A, Teeven L, Derakhshan S, Korving J, Begthel H, Dekkers JF, Kumawat K, Ramos E, van Oosterhout MF, Offerhaus GJ, Wiener DJ, Olimpio EP, Dijkstra KK, Smit EF, van der Linden M, Jaksani S, van de Ven M, Jonkers J, Rios AC, Voest EE, van Moorsel CH, van der Ent CK, Cuppen E, van Oudenaarden A, Coenjaerts FE, Meyaard L, Bont LJ, Peters PJ, Tans SJ, van Zon JS, Boj SF, Vries RG, Beekman JM, Clevers H. 2019. Long-term expanding human airway organoids for disease modeling. EMBO J 38. 10.15252/embj.2018100300.

24. Rajan A, Weaver AM, Aloisio GM, Jelinski J, Johnson HL, Venable SF, McBride T, Aideyan L, Piedra FA, Ye X, Melicoff-Portillo E, Yerramilli MRK, Zeng XL, Mancini MA, Stossi F, Maresso AW, Kotkar SA, Estes MK, Blutt S, Avadhanula V, Piedra PA. 2022. The Human Nose Organoid Respiratory Virus Model: an Ex Vivo Human Challenge Model To Study Respiratory Syncytial Virus (RSV) and Severe Acute Respiratory Syndrome Coronavirus 2 (SARS-CoV-2) Pathogenesis and Evaluate Therapeutics. mBio 13:e0351121. 10.1128/mbio.03511-21.

25. Aloisio GM, Nagaraj D, Murray AM, Schultz EM, McBride T, Aideyan L, Nicholson EG, Henke D, Ferlic-Stark L, Rajan A, Kambal A, Johnson HL, Mosa E, Stossi F, Blutt SE, Piedra PA, Avadhanula V. 2024. Infant-derived human nasal organoids exhibit relatively increased susceptibility, epithelial responses, and cytotoxicity during RSV infection. J Infect 89:106305. 10.1016/j.jinf.2024.106305.

26. Li C, Yu Y, Wan Z, Chiu MC, Huang J, Zhang S, Zhu X, Lan Q, Deng Y, Zhou Y, Xue W, Yue M, Cai JP, Yip CC, Wong KK, Liu X, Yu Y, Huang L, Chu H, Chan JF, Clevers H, Yuen KY, Zhou J. 2024. Human respiratory organoids sustained reproducible propagation of human rhinovirus C and elucidation of virus-host interaction. Nat Commun 15:10772. 10.1038/s41467-024-55076-2.

27. Zhang X, Lam SJ, Ip JD, Fong CH, Chu AW, Chan WM, Lai YS, Tsoi HW, Chan BP, Chen LL, Meng X, Yuan S, Zhao H, Cheng VC, Yuen JKY, Yuen KY, Zhou J, To KK. 2024. Characterizing fitness and immune escape of SARS-CoV-2 EG.5 sublineage using elderly serum and nasal organoid. iScience 27:109706. 10.1016/j.isci.2024.109706.

28. Bukowy-Bieryllo Z. 2021. Long-term differentiating primary human airway epithelial cell cultures: how far are we? Cell Commun Signal 19:63. 10.1186/s12964-021-00740-z.

29. Martens K, Hellings PW, Steelant B. 2018. Calu-3 epithelial cells exhibit different immune and epithelial barrier responses from freshly isolated primary nasal epithelial cells in vitro. Clin Transl Allergy 8:40. 10.1186/s13601-018-0225-8.

30. Silva S, Bicker J, Falcao A, Fortuna A. 2023. Air-liquid interface (ALI) impact on different respiratory cell cultures. Eur J Pharm Biopharm 184:62–82. 10.1016/j.ejpb.2023.01.013.

31. Kiedrowski MR, Paharik AE, Ackermann LW, Shelton AU, Singh SB, Starner TD, Horswill AR. 2016. Development of an in vitro colonization model to investigate *Staphylococcus aureus* interactions with airway epithelia. Cell Microbiol 18:720–732. 10.1111/cmi.12543.

32. De Rudder C, Calatayud Arroyo M, Lebeer S, Van de Wiele T. 2020. Dual and Triple Epithelial Coculture Model Systems with Donor-Derived Microbiota and THP-1 Macrophages To Mimic Host-Microbe Interactions in the Human Sinonasal Cavities. mSphere 5. 10.1128/mSphere.00916-19.

33. Huffines JT, Boone RL, Kiedrowski MR. 2024. Temperature influences commensal-pathogen dynamics in a nasal epithelial cell co-culture model. mSphere 9:e0058923. 10.1128/msphere.00589-23.

34. Charles DD, Fisher JR, Hoskinson SM, Medina-Colorado AA, Shen YC, Chaaban MR, Widen SG, Eaves-Pyles TD, Maxwell CA, Miller AL, Popov VL, Pyles RB. 2019. Development of a Novel ex vivo Nasal Epithelial Cell Model Supporting Colonization With Human Nasal Microbiota. Front Cell Infect Microbiol 9:165. 10.3389/fcimb.2019.00165.

35. Kokai-Kun JF, Walsh SM, Chanturiya T, Mond JJ. 2003. Lysostaphin cream eradicates *Staphylococcus aureus* nasal colonization in a cotton rat model. Antimicrob Agents Chemother 47:1589–1597. 10.1128/AAC.47.5.1589-1597.2003.

36. Bogaert D, Weinberger D, Thompson C, Lipsitch M, Malley R. 2009. Impaired innate and adaptive immunity to Streptococcus pneumoniae and its effect on colonization in an infant mouse model. Infect Immun 77:1613–1622. 10.1128/IAI.00871-08.

37. Park JC, Im SH. 2020. Of men in mice: the development and application of a humanized gnotobiotic mouse model for microbiome therapeutics. Exp Mol Med 52:1383–1396. 10.1038/s12276-020-0473-2.

38. Rajan A, Robertson MJ, Carter HE, Poole NM, Clark JR, Green SI, Criss ZK, Zhao B, Karandikar U, Xing Y, Margalef-Catala M, Jain N, Wilson RL, Bai F, Hyser JM, Petrosino J, Shroyer NF, Blutt SE, Coarfa C, Song X, Prasad BV, Amieva MR, Grande-Allen J, Estes MK, Okhuysen PC, Maresso AW. 2020. Enteroaggregative *E. coli* Adherence to Human Heparan Sulfate Proteoglycans Drives Segment and Host Specific Responses to Infection. PLoS Pathog 16:e1008851. 10.1371/journal.ppat.1008851.

39. Green SI, Gu Liu C, Yu X, Gibson S, Salmen W, Rajan A, Carter HE, Clark JR, Song X, Ramig RF, Trautner BW, Kaplan HB, Maresso AW. 2021. Targeting of Mammalian Glycans Enhances Phage Predation in the Gastrointestinal Tract. mBio 12. 10.1128/mBio.03474-20.

40. Co JY, Crouzier T, Ribbeck K. 2015. Probing the Role of Mucin-Bound Glycans in Bacterial Repulsion by Mucin Coatings. Advanced Materials Interfaces 2:1500179. 10.1002/admi.201500179.

41. Fekete E, Buret AG. 2023. The role of mucin O-glycans in microbiota dysbiosis, intestinal homeostasis, and host-pathogen interactions. Am J Physiol Gastrointest Liver Physiol 324:G452–G465. 10.1152/ajpgi.00261.2022.

42. Keck T, Leiacker R, Heinrich A, Kuhnemann S, Rettinger G. 2000. Humidity and temperature profile in the nasal cavity. Rhinology 38:167–171.

43. Bastock RA, Marino EC, Wiemels RE, Holzschu DL, Keogh RA, Zapf RL, Murphy ER, Carroll RK. 2021. *Staphylococcus aureus* Responds to Physiologically Relevant Temperature Changes by Altering Its Global Transcript and Protein Profile. mSphere 6. 10.1128/mSphere.01303-20.

44. Briaud P, Frey A, Marino EC, Bastock RA, Zielinski RE, Wiemels RE, Keogh RA, Murphy ER, Shaw LN, Carroll RK. 2021. Temperature Influences the Composition and Cytotoxicity of Extracellular Vesicles in *Staphylococcus aureus*. mSphere 6:e0067621. 10.1128/mSphere.00676-21.

45. Costa FG, Mills KB, Crosby HA, Horswill AR. 2024. The *Staphylococcus aureus* regulatory program in a human skin-like environment. mBio 15:e0045324. 10.1128/mbio.00453-24.

46. Gazioglu O, Kareem BO, Afzal M, Shafeeq S, Kuipers OP, Ulijasz AT, Andrew PW, Yesilkaya H. 2021. Glutamate Dehydrogenase (GdhA) of *Streptococcus pneumoniae* Is Required for High Temperature Adaptation. Infect Immun 89:e0040021. 10.1128/IAI.00400-21.

47. Basset A, Herd M, Daly R, Dove SL, Malley R. 2017. The Pneumococcal Type 1 Pilus Genes Are Thermoregulated and Are Repressed by a Member of the Snf2 Protein Family. J Bacteriol 199. 10.1128/JB.00078-17.

48. DeLeo FR, Otto M, Kreiswirth BN, Chambers HF. 2010. Community-associated meticillin-resistant Staphylococcus aureus. Lancet 375:1557–1568. 10.1016/S0140-6736(09)61999-1.

49. Vadia S, Tse JL, Lucena R, Yang Z, Kellogg DR, Wang JD, Levin PA. 2017. Fatty Acid Availability Sets Cell Envelope Capacity and Dictates Microbial Cell Size. Curr Biol 27:1757–1767 e1755. 10.1016/j.cub.2017.05.076.

50. Sanford BA, Thomas VL, Ramsay MA. 1989. Binding of staphylococci to mucus in vivo and in vitro. Infect Immun 57:3735–3742. 10.1128/iai.57.12.3735-3742.1989.

51. Feldman C, Read R, Rutman A, Jeffery PK, Brain A, Lund V, Mitchell TJ, Andrew PW, Boulnois GJ, Todd HC, et al. 1992. The interaction of *Streptococcus pneumoniae* with intact human respiratory mucosa *in vitro*. Eur Respir J 5:576–583.

52. Riss T, Niles A, Moravec R, Karassina N, Vidugiriene J. 2004. Cytotoxicity Assays: In Vitro Methods to Measure Dead Cells. *In* Markossian S, Grossman A, Arkin M, Auld D, Austin C, Baell J, Brimacombe K, Chung TDY, Coussens NP, Dahlin JL, Devanarayan V, Foley TL, Glicksman M, Gorshkov K, Haas JV, Hall MD, Hoare S, Inglese J, Iversen PW, Lal-Nag M, Li Z, Manro JR, McGee J, McManus O, Pearson M, Riss T, Saradjian P, Sittampalam GS, Tarselli M, Trask OJ, Jr., Weidner JR, Wildey MJ, Wilson K, Xia M, Xu X (ed.), Assay Guidance Manual, Bethesda (MD).

53. Fogh J, Fogh JM, Orfeo T. 1977. One hundred and twenty-seven cultured human tumor cell lines producing tumors in nude mice. J Natl Cancer Inst 59:221–226. 10.1093/jnci/59.1.221.

54. Moore GE, Sandberg AA. 1964. Studies of a Human Tumor Cell Line with a Diploid Karyotype. Cancer 17:170–175. 10.1002/1097-0142(196402)17:2<170::aid-cncr2820170206>3.0.co;2-n.

55. Kreft ME, Jerman UD, Lasic E, Hevir-Kene N, Rizner TL, Peternel L, Kristan K. 2015. The characterization of the human cell line Calu-3 under different culture conditions and its use as an optimized in vitro model to investigate bronchial epithelial function. Eur J Pharm Sci 69:1–9. 10.1016/j.ejps.2014.12.017.

56. Sanchez-Guzman D, Boland S, Brookes O, Mc Cord C, Lai Kuen R, Sirri V, Baeza Squiban A, Devineau S. 2021. Long-term evolution of the epithelial cell secretome in preclinical 3D models of the human bronchial epithelium. Sci Rep 11:6621. 10.1038/s41598-021-86037-0.

57. Gerber W, Svitina H, Steyn D, Peterson B, Kotze A, Weldon C, Hamman JH. 2022. Comparison of RPMI 2650 cell layers and excised sheep nasal epithelial tissues in terms of nasal drug delivery and immunocytochemistry properties. J Pharmacol Toxicol Methods 113:107131. 10.1016/j.vascn.2021.107131.

58. Martin J, Rittersberger R, Treitler S, Kopp P, Ibraimi A, Koslowski G, Sickinger M, Dabbars A, Schindowski K. 2024. Characterization of a primary cellular airway model for inhalative drug delivery in comparison with the established permanent cell lines CaLu3 and RPMI 2650. In Vitro Model 3:183–203. 10.1007/s44164-024-00079-y.

59. Correa W, Brandenburg J, Behrends J, Heinbockel L, Reiling N, Paulowski L, Schwudke D, Stephan K, Martinez-de-Tejada G, Brandenburg K, Gutsmann T. 2019. Inactivation of Bacteria by gamma-Irradiation to Investigate the Interaction with Antimicrobial Peptides. Biophys J 117:1805–1819. 10.1016/j.bpj.2019.10.012.

60. Jorgensen I, Zhang Y, Krantz BA, Miao EA. 2016. Pyroptosis triggers pore-induced intracellular traps (PITs) that capture bacteria and lead to their clearance by efferocytosis. J Exp Med 213:2113–2128. 10.1084/jem.20151613.

61. Dai Y, Zhou J, Shi C. 2023. Inflammasome: structure, biological functions, and therapeutic targets. MedComm (2020) 4:e391. 10.1002/mco2.391.

62. Di Paolo NC, Shayakhmetov DM. 2016. Interleukin 1alpha and the inflammatory process. Nat Immunol 17:906–913. 10.1038/ni.3503.

63. Kim B, Lee Y, Kim E, Kwak A, Ryoo S, Bae SH, Azam T, Kim S, Dinarello CA. 2013. The Interleukin-1alpha Precursor is Biologically Active and is Likely a Key Alarmin in the IL-1 Family of Cytokines. Front Immunol 4:391. 10.3389/fimmu.2013.00391.

64. Cole AL, Muthukrishnan G, Chong C, Beavis A, Eade CR, Wood MP, Deichen MG, Cole AM. 2016. Host innate inflammatory factors and staphylococcal protein A influence the duration of human *Staphylococcus aureus* nasal carriage. Mucosal Immunol 9:1537–1548. 10.1038/mi.2016.2.

65. Liu M, Guo S, Hibbert JM, Jain V, Singh N, Wilson NO, Stiles JK. 2011. CXCL10/IP-10 in infectious diseases pathogenesis and potential therapeutic implications. Cytokine Growth Factor Rev 22:121–130. 10.1016/j.cytogfr.2011.06.001.

66. Crawford MA, Ward AE, Gray V, Bailer P, Fisher DJ, Kubicka E, Cui Z, Luo Q, Gray MC, Criss AK, Lum LG, Tamm LK, Letteri RA, Hughes MA. 2023. Disparate Regions of the Human Chemokine CXCL10 Exhibit Broad-Spectrum Antimicrobial Activity against Biodefense and Antibiotic-Resistant Bacterial Pathogens. ACS Infect Dis 9:122–139. 10.1021/acsinfecdis.2c00456.

67. Yung SC, Parenti D, Murphy PM. 2011. Host chemokines bind to *Staphylococcus aureus* and stimulate protein A release. J Biol Chem 286:5069–5077. 10.1074/jbc.M110.195180.

68. Lore NI, De Lorenzo R, Rancoita PMV, Cugnata F, Agresti A, Benedetti F, Bianchi ME, Bonini C, Capobianco A, Conte C, Corti A, Furlan R, Mantegani P, Maugeri N, Sciorati C, Saliu F, Silvestri L, Tresoldi C, Bio Angels for C-BSG, Ciceri F, Rovere-Querini P, Di Serio C, Cirillo DM, Manfredi AA. 2021. CXCL10 levels at hospital admission predict COVID-19 outcome: hierarchical assessment of 53 putative inflammatory biomarkers in an observational study. Mol Med 27:129. 10.1186/s10020-021-00390-4.

69. Klingler AI, Stevens WW, Tan BK, Peters AT, Poposki JA, Grammer LC, Welch KC, Smith SS, Conley DB, Kern RC, Schleimer RP, Kato A. 2021. Mechanisms and biomarkers of inflammatory endotypes in chronic rhinosinusitis without nasal polyps. J Allergy Clin Immunol 147:1306–1317. 10.1016/j.jaci.2020.11.037.

70. Nicholson EG, Schlegel C, Garofalo RP, Mehta R, Scheffler M, Mei M, Piedra PA. 2016. Robust Cytokine and Chemokine Response in Nasopharyngeal Secretions: Association With Decreased Severity in Children With Physician Diagnosed Bronchiolitis. J Infect Dis 214:649–655. 10.1093/infdis/jiw191.

71. Hurst JH, McCumber AW, Aquino JN, Rodriguez J, Heston SM, Lugo DJ, Rotta AT, Turner NA, Pfeiffer TS, Gurley TC, Moody MA, Denny TN, Rawls JF, Clark JS, Woods CW, Kelly MS. 2022. Age-Related Changes in the Nasopharyngeal Microbiome Are Associated With Severe Acute Respiratory Syndrome Coronavirus 2 (SARS-CoV-2) Infection and Symptoms Among Children, Adolescents, and Young Adults. Clin Infect Dis 75:e928–e937. 10.1093/cid/ciac184.

72. Li Z, Levast B, Madrenas J. 2017. *Staphylococcus aureus* Downregulates IP-10 Production and Prevents Th1 Cell Recruitment. J Immunol 198:1865–1874. 10.4049/jimmunol.1601336.

73. Urban BC, Goncalves ANA, Loukov D, Passos FM, Reine J, Gonzalez-Dias P, Solorzano C, Mitsi E, Nikolaou E, O’Connor D, Collins AM, Adler H, Pollard A, Rylance J, Gordon SB, Jochems SP, Nakaya HI, Ferreira DM. 2024. Inflammation of the nasal mucosa is associated with susceptibility to experimental pneumococcal challenge in older adults. Mucosal Immunol. 10.1016/j.mucimm.2024.06.010.

74. Tokunaga R, Zhang W, Naseem M, Puccini A, Berger MD, Soni S, McSkane M, Baba H, Lenz HJ. 2018. CXCL9, CXCL10, CXCL11/CXCR3 axis for immune activation - A target for novel cancer therapy. Cancer Treat Rev 63:40–47. 10.1016/j.ctrv.2017.11.007.

75. Seyoum B, Yano M, Pirofski LA. 2011. The innate immune response to *Streptococcus pneumoniae* in the lung depends on serotype and host response. Vaccine 29:8002–8011. 10.1016/j.vaccine.2011.08.064.

76. Hieshima K, Imai T, Opdenakker G, Van Damme J, Kusuda J, Tei H, Sakaki Y, Takatsuki K, Miura R, Yoshie O, Nomiyama H. 1997. Molecular cloning of a novel human CC chemokine liver and activation-regulated chemokine (LARC) expressed in liver. Chemotactic activity for lymphocytes and gene localization on chromosome 2. J Biol Chem 272:5846–5853. 10.1074/jbc.272.9.5846.

77. Le Borgne M, Etchart N, Goubier A, Lira SA, Sirard JC, van Rooijen N, Caux C, Ait-Yahia S, Vicari A, Kaiserlian D, Dubois B. 2006. Dendritic cells rapidly recruited into epithelial tissues via CCR6/CCL20 are responsible for CD8+ T cell crosspriming in vivo. Immunity 24:191–201. 10.1016/j.immuni.2006.01.005.

78. Martin KR, Wong HL, Witko-Sarsat V, Wicks IP. 2021. G-CSF - A double edge sword in neutrophil mediated immunity. Semin Immunol 54:101516. 10.1016/j.smim.2021.101516.

79. Cantero D, Cooksley C, Jardeleza C, Bassiouni A, Jones D, Wormald PJ, Vreugde S. 2013. A human nasal explant model to study *Staphylococcus aureus* biofilm *in vitro*. Int Forum Allergy Rhinol 3:556–562. 10.1002/alr.21146.

80. Hu H, Liu S, Hon K, Psaltis AJ, Wormald PJ, Vreugde S. 2023. Staphylococcal protein A modulates inflammation by inducing interferon signaling in human nasal epithelial cells. Inflamm Res 72:251–262. 10.1007/s00011-022-01656-1.

81. McNab F, Mayer-Barber K, Sher A, Wack A, O’Garra A. 2015. Type I interferons in infectious disease. Nat Rev Immunol 15:87–103. 10.1038/nri3787.

82. Kovarik P, Castiglia V, Ivin M, Ebner F. 2016. Type I Interferons in Bacterial Infections: A Balancing Act. Front Immunol 7:652. 10.3389/fimmu.2016.00652.

83. Rodenburg LW, Metzemaekers M, van der Windt IS, Smits SMA, den Hertog-Oosterhoff LA, Kruisselbrink E, Brunsveld JE, Michel S, de Winter-de Groot KM, van der Ent CK, Stadhouders R, Beekman JM, Amatngalim GD. 2023. Exploring intrinsic variability between cultured nasal and bronchial epithelia in cystic fibrosis. Sci Rep 13:18573. 10.1038/s41598-023-45201-4.

84. Pleguezuelos-Manzano C, Beenker WAG, van Son GJF, Begthel H, Amatngalim GD, Beekman JM, Clevers H, den Hertog J. 2025. Dual RNA sequencing of a co-culture model of Pseudomonas aeruginosa and human 2D upper airway organoids. Sci Rep 15:2222. 10.1038/s41598-024-82500-w.

85. Chiu MC, Li C, Liu X, Song W, Wan Z, Yu Y, Huang J, Xiao D, Chu H, Cai JP, To KK, Yuen KY, Zhou J. 2022. Human Nasal Organoids Model SARS-CoV-2 Upper Respiratory Infection and Recapitulate the Differential Infectivity of Emerging Variants. mBio 13:e0194422. 10.1128/mbio.01944-22.

86. Li L, Jiao L, Feng D, Yuan Y, Yang X, Li J, Jiang D, Chen H, Meng Q, Chen R, Fang B, Zou X, Luo Z, Ye X, Hong Y, Liu C, Li C. 2024. Human apical-out nasal organoids reveal an essential role of matrix metalloproteinases in airway epithelial differentiation. Nat Commun 15:143. 10.1038/s41467-023-44488-1.

87. Lin SC, Qu L, Ettayebi K, Crawford SE, Blutt SE, Robertson MJ, Zeng XL, Tenge VR, Ayyar BV, Karandikar UC, Yu X, Coarfa C, Atmar RL, Ramani S, Estes MK. 2020. Human norovirus exhibits strain-specific sensitivity to host interferon pathways in human intestinal enteroids. Proc Natl Acad Sci U S A 117:23782–23793. 10.1073/pnas.2010834117.

88. Zushin PH, Mukherjee S, Wu JC. 2023. FDA Modernization Act 2.0: transitioning beyond animal models with human cells, organoids, and AI/ML-based approaches. J Clin Invest 133. 10.1172/JCI175824.

89. Karp PH, Moninger TO, Weber SP, Nesselhauf TS, Launspach JL, Zabner J, Welsh MJ. 2002. An in vitro model of differentiated human airway epithelia. Methods for establishing primary cultures. Methods Mol Biol 188:115–137. 10.1385/1-59259-185-X:115.

90. Srinivasan B, Kolli AR, Esch MB, Abaci HE, Shuler ML, Hickman JJ. 2015. TEER measurement techniques for in vitro barrier model systems. J Lab Autom 20:107–126. 10.1177/2211068214561025.

91. Liao Y, Smyth GK, Shi W. 2014. featureCounts: an efficient general purpose program for assigning sequence reads to genomic features. Bioinformatics 30:923–930. 10.1093/bioinformatics/btt656.

92. Love MI, Huber W, Anders S. 2014. Moderated estimation of fold change and dispersion for RNA-seq data with DESeq2. Genome Biol 15:550. 10.1186/s13059-014-0550-8.

93. Flores Ramos S, Brugger SD, Escapa IF, Skeete CA, Cotton SL, Eslami SM, Gao W, Bomar L, Tran TH, Jones DS, Minot S, Roberts RJ, Johnston CD, Lemon KP. 2021. Genomic Stability and Genetic Defense Systems in *Dolosigranulum pigrum*, a Candidate Beneficial Bacterium from the Human Microbiome. mSystems 6:e0042521. 10.1128/mSystems.00425-21.

94. Fey PD, Endres JL, Yajjala VK, Widhelm TJ, Boissy RJ, Bose JL, Bayles KW. 2013. A genetic resource for rapid and comprehensive phenotype screening of nonessential *Staphylococcus aureus* genes. mBio 4:e00537–00512. 10.1128/mBio.00537-12.

95. Malley R, Lipsitch M, Stack A, Saladino R, Fleisher G, Pelton S, Thompson C, Briles D, Anderson P. 2001. Intranasal immunization with killed unencapsulated whole cells prevents colonization and invasive disease by capsulated pneumococci. Infect Immun 69:4870–4873. 10.1128/IAI.69.8.4870-4873.2001.

96. R-Core-Team. 2021. R: A Language and Environment for Statistical Computing. R Foundation for Statistical Computing, Vienna, Austria.

97. RStudio-Team. 2020. RStudio: Integrated Development for R. RStudio. PBC, Boston, MA.

98. Bates D, Mächler M, Bolker B, Walker S. 2015. Fitting Linear Mixed-Effects Models Using lme4. Journal of Statistical Software 67:1–48. 10.18637/jss.v067.i01.

99. Holm S. 1979. A Simple Sequentially Rejective Multiple Test Procedure. Scandinavian Journal of Statistics 6:65–70.

100. Singmann H, Bolker, B., Westfall, J., Aust, F., Ben-Shachar, M. S. 2024. afex: Analysis of Factorial Experiments.

101. Lenth RV. 2024. emmeans: Estimated Marginal Means, aka Least-Squares Means.

102. Benjamini Y, Hochberg Y. 1995. Controlling the False Discovery Rate: A Practical and Powerful Approach to Multiple Testing. Journal of the Royal Statistical Society. Series B (Methodological) 57:289–300.

